# Iron as a principal mediator of dysbiosis prevention in the phyllosphere

**DOI:** 10.64898/2026.09.16.751549

**Authors:** Shanice S. Webster, Jin Xu, Mao Kobayashi, Jingyuan Zhang, Tao Chen, Sheng Yang He

**Author notes:** Correspondence (S.S.W.) and (S.Y.H.).

## Abstract

Multicellular organisms require a correct microbiota composition for optimal health and fitness. Dysbiosis could lead to serious health consequences in humans and plants, and is pervasive during pathogen infections across systems. A recent study shows that carbon source availability plays a role in bacterial community assembly in *Arabidopsis* leaves [1]. However, whether carbon sources or other molecules are critical for leaf microbiota homeostasis remains unknown. Here, an unbiased *in planta* metatranscriptomic analysis of endophytic microbiota in leaves of wild-type and dysbiotic *min7 fls2 efr cerk1* (*mfec*) mutant, which is defective in pattern-recognition receptor (PRR) signaling and MIN7-dependent vesicle trafficking, revealed that bacteria inside *Arabidopsis* leaves exhibit taxa-specific transcriptional responses. Most notably, iron starvation response was detected in *Stenotrophomonas* strains, a group of endophytic bacteria that cause dysbiotic symptoms. Elevated expression of canonical iron starvation genes confirmed low iron availability in *mfec* plants. This low iron environment was associated with a reduced level of the plasma membrane H -ATPase AHA2, resulting in a higher apoplastic pH that favors less-bioavailable ferric iron over soluble ferrous iron. Supplementing iron in *mfec* plants was sufficient to restore microbiota homeostasis and alleviate dysbiosis phenotypes. The role of iron in dysbiosis extends to *Pseudomonas syringae* foliar infection, which drives leaf dysbiosis through two iron-dependent mechanisms. Iron thus emerges as a surprising, key switch between healthy and dysbiotic microbiota in the endophytic spaces of plants.

## INTRODUCTION

Microbial communities are integral to the health and function of nearly all multicellular organisms. In humans, dysbiosis contributes to a variety of illnesses [2, 3]. In plants, dysbiosis is associated with severe tissue damage and/or autoimmunity [4–6]. How plants and humans prevent dysbiosis is a fundamentally important but not a well understood question. The aboveground parts of plants harbor some of the most diverse microbial communities on Earth [7]. These microbial communities inhabit both the leaf surfaces (epiphytic microbiota) and the leaf interior (endophytic microbiota) and are vital for plant growth, stress tolerance, and protection against pathogens [8]. Despite constant exposure to a large pool of soil-derived microbes during germination and airborne microbes thereafter, only a small subset of microbes successfully colonize the leaf interior, indicating that plants actively regulate microbiota abundance and composition within this endophytic environment. We previously discovered that dysbiosis in leaves is associated with an over-proliferation and drastically altered microbiota composition, characterized by an increase of certain Proteobacteria members, such as *Stenotrophomonas* strains, and depletion of Firmicutes, such as *Paenibacillus* strains [5]. Little is known, however, about the specific plant-derived metabolic signals, and how their levels determine whether the endophytic community remains healthy or shifts into dysbiosis. Understanding how plants tune these signals to the correct levels is central to explaining why endophytic communities remain compositionally stable under normal conditions yet can collapse into dysbiosis when this balance is disrupted. Of note, dysbiosis is a recurring feature of pathogen and pest infestation across diverse plant-microbe and plant-insect systems [9, 10].

Progress has been made to elucidate host mechanisms that are involved in shaping a healthy microbiota. Hosts across many systems maintain stable symbiotic communities by actively controlling the nutrients and signals available to colonizing microbes. A recent study of 224 bacterial strains from *Arabidopsis* leaves found that carbon source availability and metabolic interactions among strains shape bacterial community assembly in the leaf [1]. On the other hand, genome-wide association and mutational studies of plants, have implicated genes involved in immune signaling, vesicle trafficking, MACPF-family protein, S-acyltransferase, cell wall remodeling in controlling leaf microbiota [5, 11–14]. In particular, the *Arabidopsis mfec* quadruple mutant, with dual defects in PRR signaling and MIN7-dependent vesicle trafficking, causally exhibits spontaneous tissue-damaging dysbiosis in leaves [5]. Despite these advances, the metabolic basis of the switch between a healthy and a dysbiotic leaf microbiota remains undefined.

In this study, we used *in planta* metatranscriptomics to directly probe the endophytic leaf environments of wild-type Col-0 and the *mfec* plants. We identify iron as a key factor governing the switch between healthy and dysbiotic microbiota in *Arabidopsis* leaves. This finding has significant basic and applied implications in dysbiosis prevention.

## RESULTS

### *S. maltophilia* (C46) transcriptome reveals an iron-limited endophytic environment in *mfec* leaves

To probe the endophytic metabolic environment of the leaf, we used *S. maltophilia* (C46) as a microbial sensor of Col-0 (WT) and *mfec* leaves. We chose C46 because the relative abundance of *Stenotrophomonas* strains was shown to be sensitive to host genotype, being among the most enriched taxa in the endosphere of *mfec* leaves compared to WT leaves [5]. In addition, *Stenotrophomonas* strains were the most potent strains that causally induce dysbiosis-associated tissue damage [5]. We infiltrated C46 alone into WT and *mfec* leaves and compared its transcriptome between host genotypes (Fig. 1a). Principal coordinates analysis (PCoA) separated samples primarily by host genotype along PCoA1 (40.53% of variance explained), showing that C46 adopts distinct transcriptional states depending on whether it resides in WT or *mfec* leaves (Extended Data Fig. 1a). Consistent with this, differential expression analysis identified 193 genes upregulated and 268 genes downregulated in *mfec* relative to WT leaves (Extended Data Fig. 1b). Most notably, biosynthesis of siderophore group non-ribosomal peptides was the most strongly and significantly enriched pathway among genes upregulated in *mfec* leaves, indicating that C46 perceives the *mfec* endophytic environment as an iron-limited nutritional environment (Fig. 1b). Vitamin B6 metabolism was also significantly enriched among upregulated genes in *mfec* leaves, consistent with the involvement of B6 in iron starvation response [15] (Supplementary Table S1). Genes upregulated in WT relative to *mfec* leaves were strongly enriched for non-homologous end joining, plant-pathogen interactions and bacterial motility proteins (Fig. 1c). Upregulation of bacterial motility proteins was reported as a bacterial response to plant immune activation [16], which is consistent with the expectation that C46 senses a higher immune activity in WT than in *mfec* leaves.

We previously established a 48-member endophytic bacterial synthetic community (SynCom^Col-0^) from healthy WT leaves [5, 17]. To determine whether the transcriptomic features of C46 were retained when C46 is inoculated as a member of a complex microbiota, a condition that better reflects its natural context, we performed *in planta* metatranscriptomics using SynCom^Col-0^ that includes C46 (Fig. 1d). As with “C46 alone” inoculation, PCoA of the C46 transcriptome within the SynCom separated samples by host genotype, explaining 54.2% of the variance and demonstrate that genotype-dependent transcriptional divergence of C46 is retained in a community context (Extended Data Fig. 1c). This was accompanied by 76 up- and 145 downregulated C46 genes from *mfec* leaves compared to C46 from WT leaves (Extended Data Fig. 1d). As with “C46 alone” inoculation, biosynthesis of siderophore group non-ribosomal peptides in C46 remained one of the most strongly and significantly enriched pathways among genes upregulated in *mfec* leaves (Fig. 1e). Conversely, genes upregulated in WT relative to *mfec* leaves were most strongly enriched for flagellar assembly, bacterial motility/chemotaxis, alongside plant-pathogen interactions, largely mirroring the pathways enriched in WT when C46 was infiltrated into leaves alone (Fig. 1f). These results show that iron acquisition is activated in *mfec* leaves, and motility/chemotaxis behaviors are activated in WT leaves, regardless of whether C46 is inoculated into leaves alone or inoculated with a complex community. The *ent*-like biosynthetic operon in C46 (*entC*, *entE*, *entB*, *entD*, *entF*, *entA*) is homologous to the *E. coli* enterobactin pathway, which encodes a catecholate siderophore induced under iron limitation. Consistent with pathway-level enrichment, these genes showed elevated expression in *mfec* relative to WT leaves in both “C46 alone” and “C46 with other SynCom^Col-0^ strains” albeit to different degrees (Extended Data Fig. 1e, f, Supplementary Table S1 and S2).

**Figure 1.**
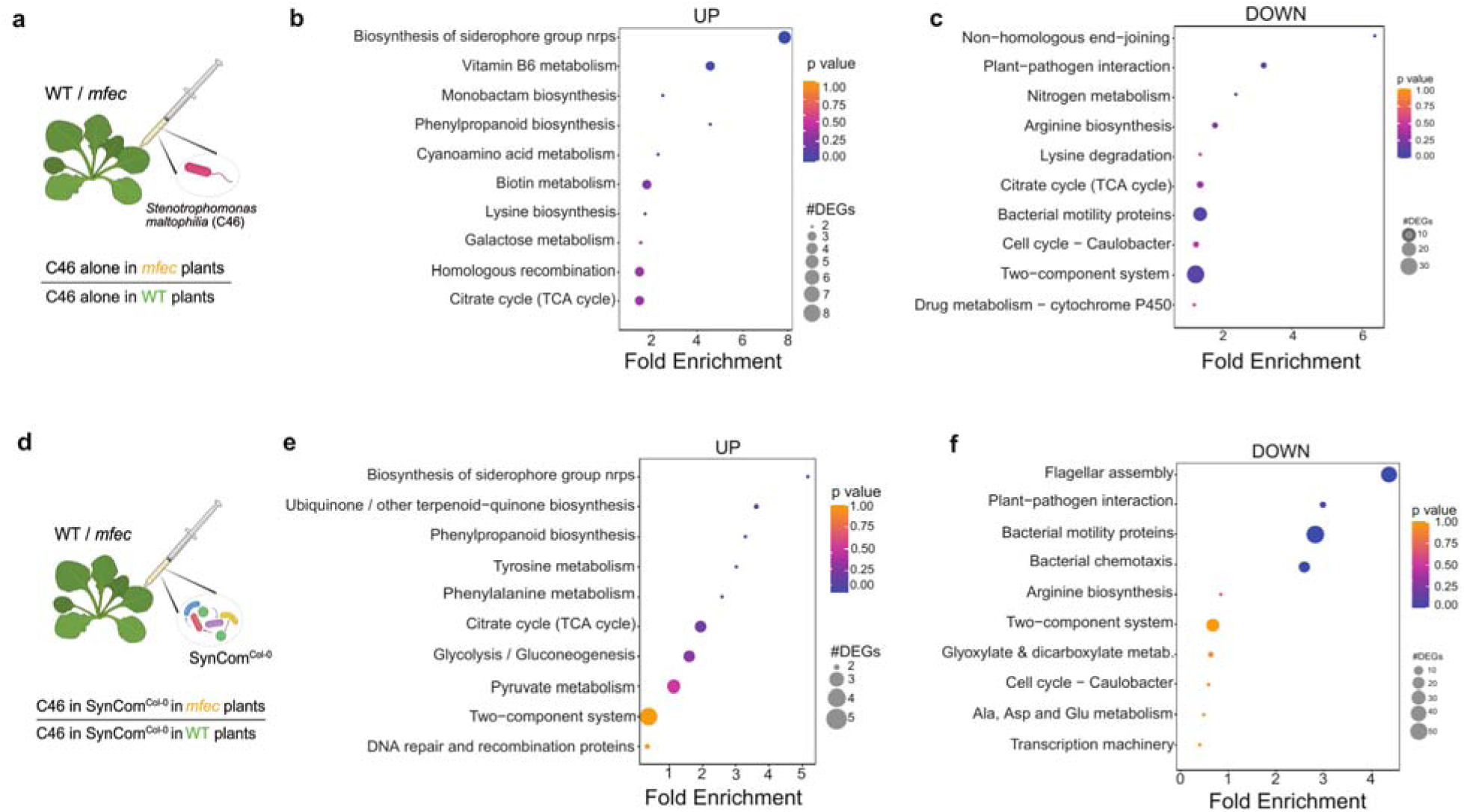
Transcriptomic response of *S. maltophilia* (C46) to host genotype and microbiota community context. **a**, Schematic of C46 mono-infiltration experiment: C46 alone was infiltrated in WT Col-0 and *mfec* leaves and bacterial RNA was extracted for RNA-sequencing (n = 4 biological replicates). C46 in WT plants was used as the reference condition in differential expression analyses. Differentially expressed genes (DEGs) were identified using DESeq2 with a threshold of (padj < 0.05). **b,c,** KEGG pathway enrichment analysis using Fisher’s exact test (p adjusted < 0.05) of genes significantly (**b**) upregulated or (**c**) downregulated in C46 in *mfec* relative to WT leaves. The top 10 pathways with the lowest adjusted p values is shown. Dot size indicates the number of DEGs in each pathway, color indicates enrichment p-value and x-axis shows fold enrichment. **d**, Schematic of C46 infiltrated as part of SynCom^Col-0^ in WT and *mfec* leaves.**e,f,** KEGG pathway enrichment analysis of genes (**e**) upregulated or (**f**) downregulated in C46 in *mfec* versus WT leaves when part of the SynCom. Dot sizes and color as in (**b,c**).

**Extended Data Fig. 1.**
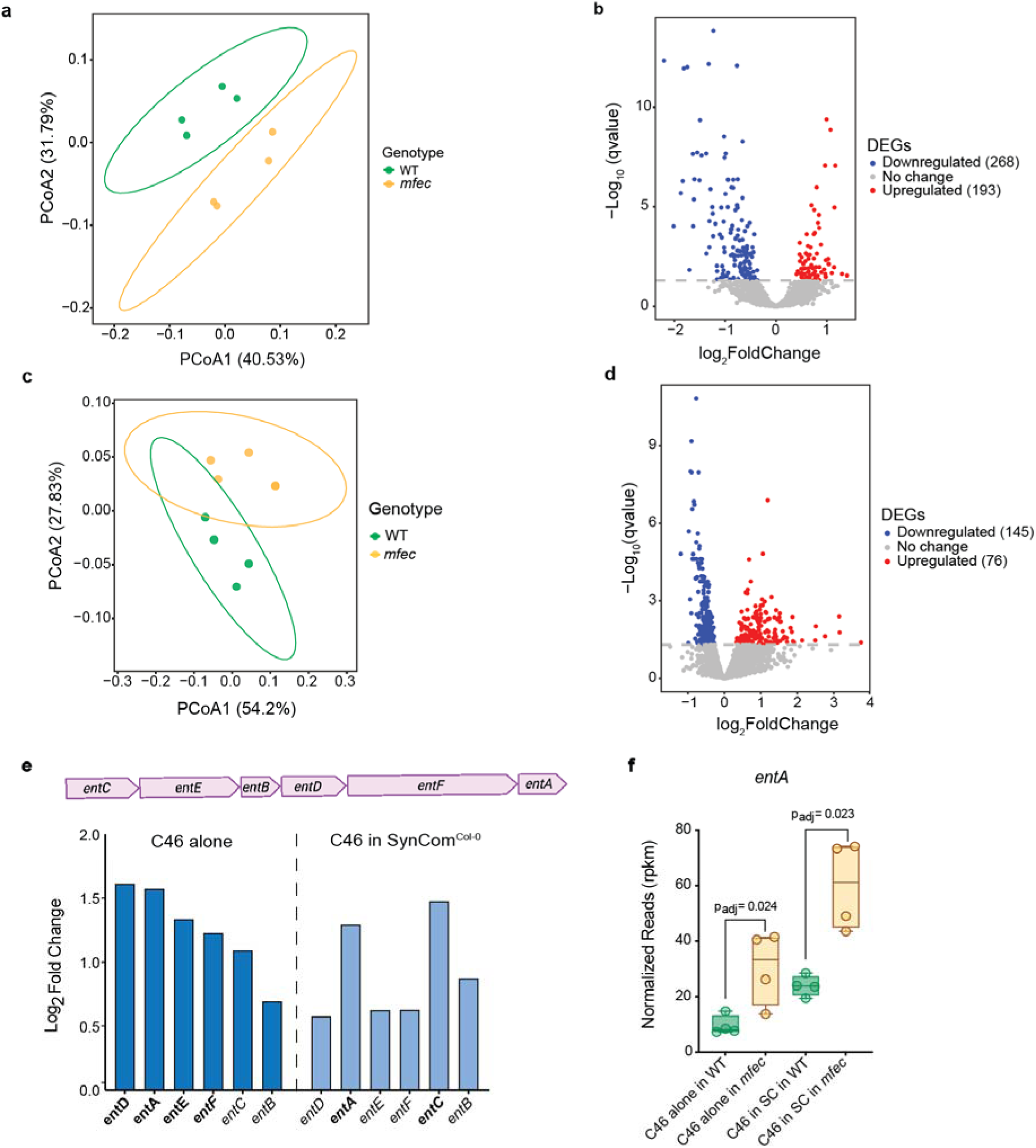
*S. maltophilia* (C46) undergoes transcriptional reprogramming including upregulation of enterobactin-family siderophore biosynthesis genes, **a**, Principal coordinates analysis (PCoA) of C46 transcriptomes following mono-infiltration into WT and *mfec* leaves, based on Bray-Curtis dissimilarity. Each point represents one biological replicate (n = 4), colored by host genotype. **b**, Volcano plot of differentially expressed genes (DEGs) in C46 alone in *mfec* versus WT leaves. Genes are colored by direction of regulation: downregulated (dark blue, n = 268), upregulated (red, n = 193), and not significantly changed (grey). Dashed line indicates the significance threshold (q = 0.05). **c**, PCoA of C46 transcriptomes when infiltrated as part of SynCom^Col-0^ into WT and *mfec* leaves. **d**, Volcano plot of DEGs in C46 within SynCom^Col-0^ in *mfec* versus WT leaves. Genes are colored by direction of regulation, as in b: downregulated (dark blue, n = 145), upregulated (red, n = 76), and not significantly changed (grey). **e,** (Top) Diagram of the enterobactin-like siderophore biosynthesis operon (*entAFDBEC*). (Bottom) Log_2_ fold change (*mfec* vs. WT) for each gene in the operon, shown separately for C46 alone (left) and C46 when with SynCom^Col-0^ (right). Bold gene names indicate genes with statistically significant differential expression (padj < 0.05). **f**, Normalized expression (reads per kilobase per million mapped reads, rpkm) of *entA* across all four experimental conditions: C46 alone in WT, C46 alone in *mfec*, C46 in SynCom^Col-0^ (SC) in WT, and C46 in SynCom^Col-0^ (SC) in *mfec*. Each dot represents one biological replicate (n = 4). Adjusted p-values (padj) were calculated using a DESeq2 Wald test with Benjamini-Hochberg correction for multiple testing.

### *mfec* plants exhibit systemic iron deficiency responses

Because upregulation of two-component system and bacterial motility proteins was reported as a bacterial response to plant immune activation [16], the downregulation of bacterial two-component system and flagellum/motility proteins in *mfec* leaves was therefore expected. However, we were surprised by the exaggerated iron deficiency transcriptomic signatures in C46 colonizing *mfec* leaves compared to WT leaves. To confirm whether *mfec* plants are indeed experiencing iron deficiency, we performed RNA sequencing of 2.5- and 4.5-week-old potting soil-grown WT and *mfec* plants (*i.e*., colonized by microbiota from potting soil, water and/or circulating air in the growth chamber) to capture the iron response status both at an early and a later stage of vegetative growth. Canonical iron-responsive genes span several functional categories, including iron deficiency signaling, iron transport and uptake, iron storage, and systemic iron signaling (Fig. 2a, Extended Data Fig. 2a-d, Supplementary Table S3, S4). Genes across all of these categories were found to be altered in *mfec* leaves, for example, iron deficiency-responsive basic helix-loop-helix (bHLH) transcription factors, including *bHLH38*, *bHLH39*, *bHLH100*, and *bHLH101*, along with the deficiency-signaling gene *BTSL1*, were increased in *mfec* plants (Fig. 2b, Extended Data Fig. 2a-d). In contrast, genes involved in iron storage, including *FER1*, *FER2* and *FER3*, showed reduced expression in *mfec* leaves, consistent with the known suppression of iron storage under iron-limiting conditions [18] (Fig. 2b, Extended Data Fig. 2d). To determine whether iron deficiency extends to the roots, we examined protein levels of the canonical iron-regulated transporter (IRT1) protein. *mfec* roots expressed higher IRT1 protein levels than WT roots (Fig. 2d), demonstrating that altered iron status is not confined to aerial tissues but is accompanied by activation of root iron acquisition pathways. Together, these data confirm that *mfec* plants systemically experience low iron availability compared to WT. This prompted us to measure the iron content in WT vs. *mfec* leaves. However, the iron content in *mfec* leaves was only slightly lower than that in WT leaves (Fig. 2e).

**Figure 2.**
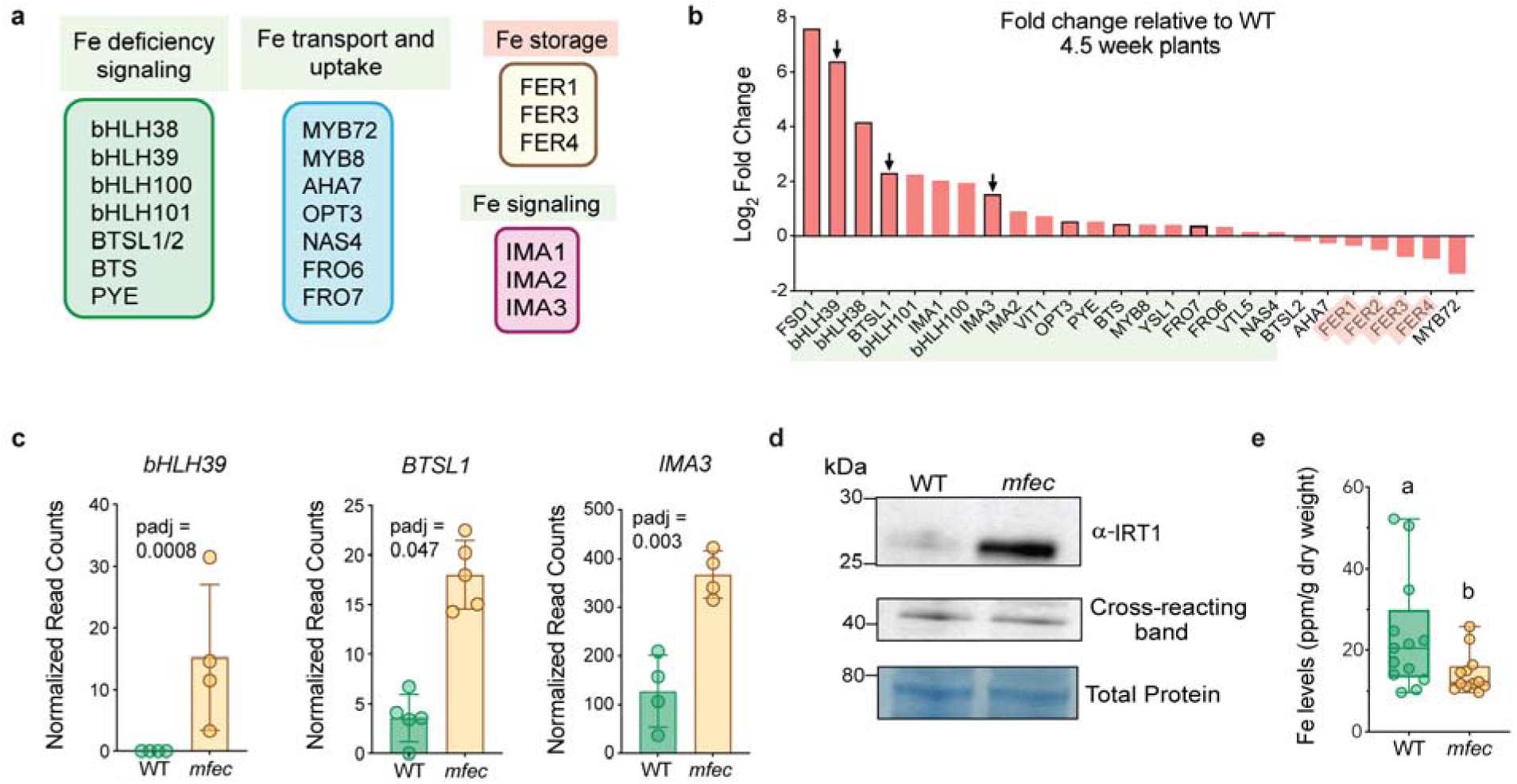
*mfec* plants show increased iron starvation at the transcript and protein level. **a**, Functional categories of canonical iron homeostasis genes examined in this study, grouped by role. **b,** Log_2_ fold change of iron homeostasis genes in 4.5-week-old *mfec* plants relative to WT ranked from most upregulated to most downregulated. Arrows indicate *bHLH39, BTSL1,* and *IMA3*, shown individually in (**c**). **c**, Normalized read counts of *bHLH39, BTSL1* and *IMA3* in WT and *mfec* leaves, from DESeq2 differential expression analysis. Each dot represents one biological replicate (n = 4); bars show mean ± SEM. Adjusted p-values (padj) from DESeq2 Wald tests with Benjamini-Hochberg correction are shown for each gene. **d,** Representative western blot of IRT1 protein levels in WT and *mfec* roots (top), with a non-specific cross-reacting band (middle) and total protein stain (bottom) as loading controls. Source data for unprocessed blot is in Supplementary Fig. 1a,b. The experiment was performed three independent times **e,** Fe levels (ppm/g dry weight) in WT and *mfec* leaf tissue. Groups not sharing a letter (**a**, **b**) are significantly different (Student’s t-test, p < 0.05). Data in (**e**) are from plants collected from three independent experiments.

**Extended Data Fig. 2.**
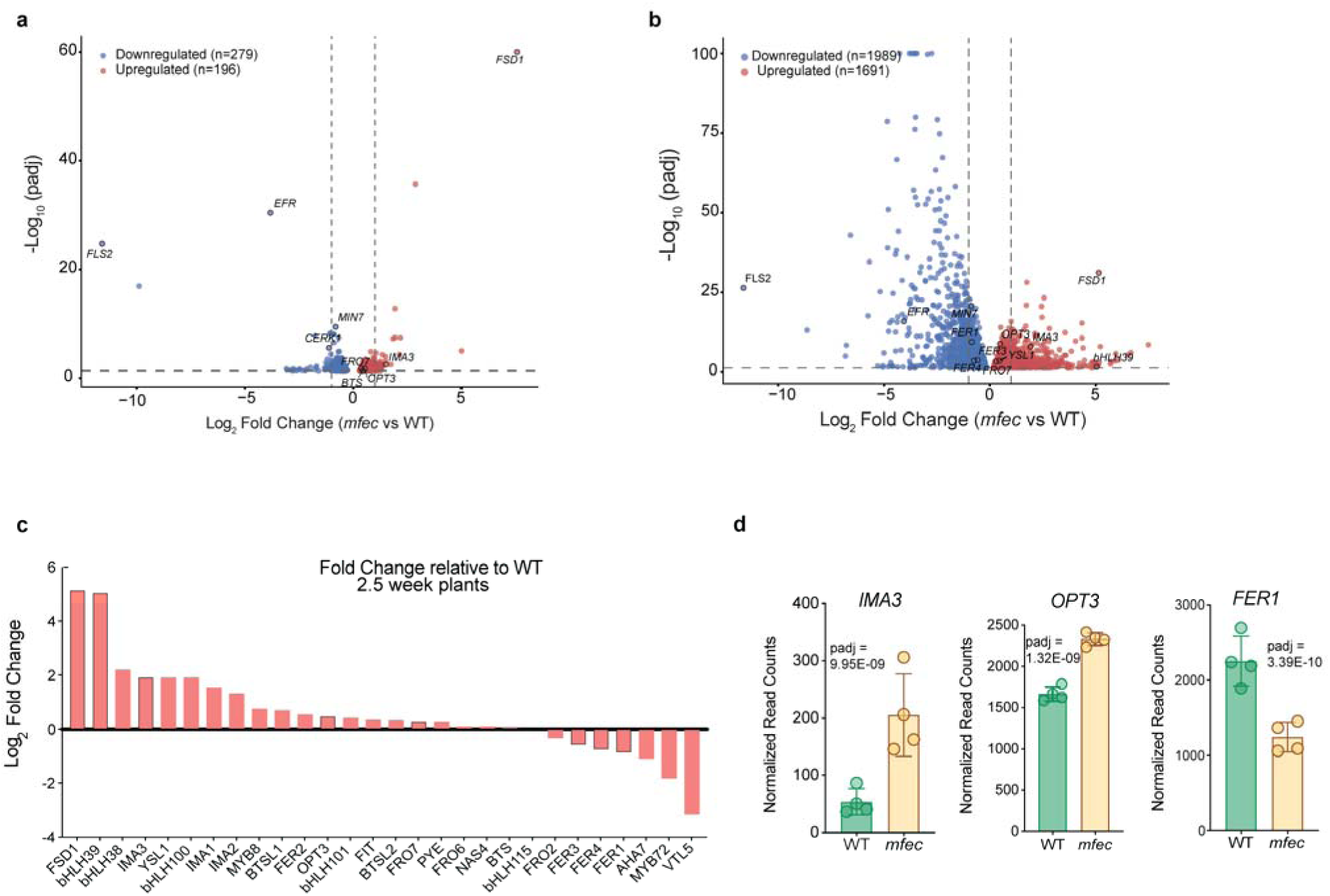
*mfec* plants exhibit iron starvation transcriptional response at 2.5 weeks. **a**, Volcano plot of differentially expressed genes (DEGs) between WT and *mfec* leaves at 4.5 weeks. Downregulated (blue, n = 279) and upregulated (red, n = 196) genes are shown relative to padj < 0.05 (dashed line) and |log_2_FC| > 1 (dotted lines). Selected genes are labeled, including *FLS2*, *EFR*, *CERK1*, and *MIN7* (disrupted in the *mfec* mutant) and canonical iron-homeostasis genes. **b**, Volcano plot of DEGs between *mfec* and WT leaves at 2.5 weeks, as in **a** (downregulated, n = 1,989; upregulated, n = 1,691). **c**, Log_2_ fold change of iron-homeostasis genes in 2.5-week-old *mfec* plants relative to WT, ranked from most upregulated to most downregulated. **d**, Normalized read counts of *IMA3*, *OPT3*, and *FER1* in WT and *mfec* leaves. Each dot represents one biological replicate (n = 4); bars represent mean ± SEM. Adjusted p-values (padj) from DESeq2 Wald test with Benjamini-Hochberg correction for multiple testing.

### The *mfec* leaf apoplast is more alkaline and correlated with AHA2 downregulation

The activation of systemic iron deficiency responses in the absence of a drastic difference in the iron content between WT and *mfec* leaves raised an apparent paradox. Iron deficiency signaling is typically induced when bioavailable iron is limiting. Because iron solubility decreases dramatically with increasing pH [19], we hypothesized that the iron starvation responses observed in *mfec* leaves might stem from reduced iron bioavailability due to apoplast alkalinization. To test this hypothesis, we measured apoplast pH using the ratiometric fluorescent pH dye 8-hydroxypyrene-1,3,6-trisulfonic acid trisodium salt (HPTS) [20] in WT and *mfec* leaves. The 405/458 (protonated to deprotonated) ratio was lower in *mfec* leaves (Fig. 3a), consistent with a more alkaline leaf apoplast than WT. Apoplastic acidification in roots and leaves is primarily mediated by the H⁺-ATPase AHA2 [21], with additional contributions from other H⁺ transporters such as NRT1.1, a dual-affinity nitrate/H⁺ transporter [22]. We therefore asked whether expression of these transporters was altered in *mfec* leaves. Transcriptomic analysis showed that *NRT1.1* was upregulated in *mfec* leaves, while *AHA2* was downregulated (Fig. 3c,d,f), indicating transcriptional dysregulation of both genes. Consistent with the RNA-seq data, immunoblot analysis showed that total AHA2 protein was also reduced in *mfec* leaves (Fig. 3e).

**Figure 3.**
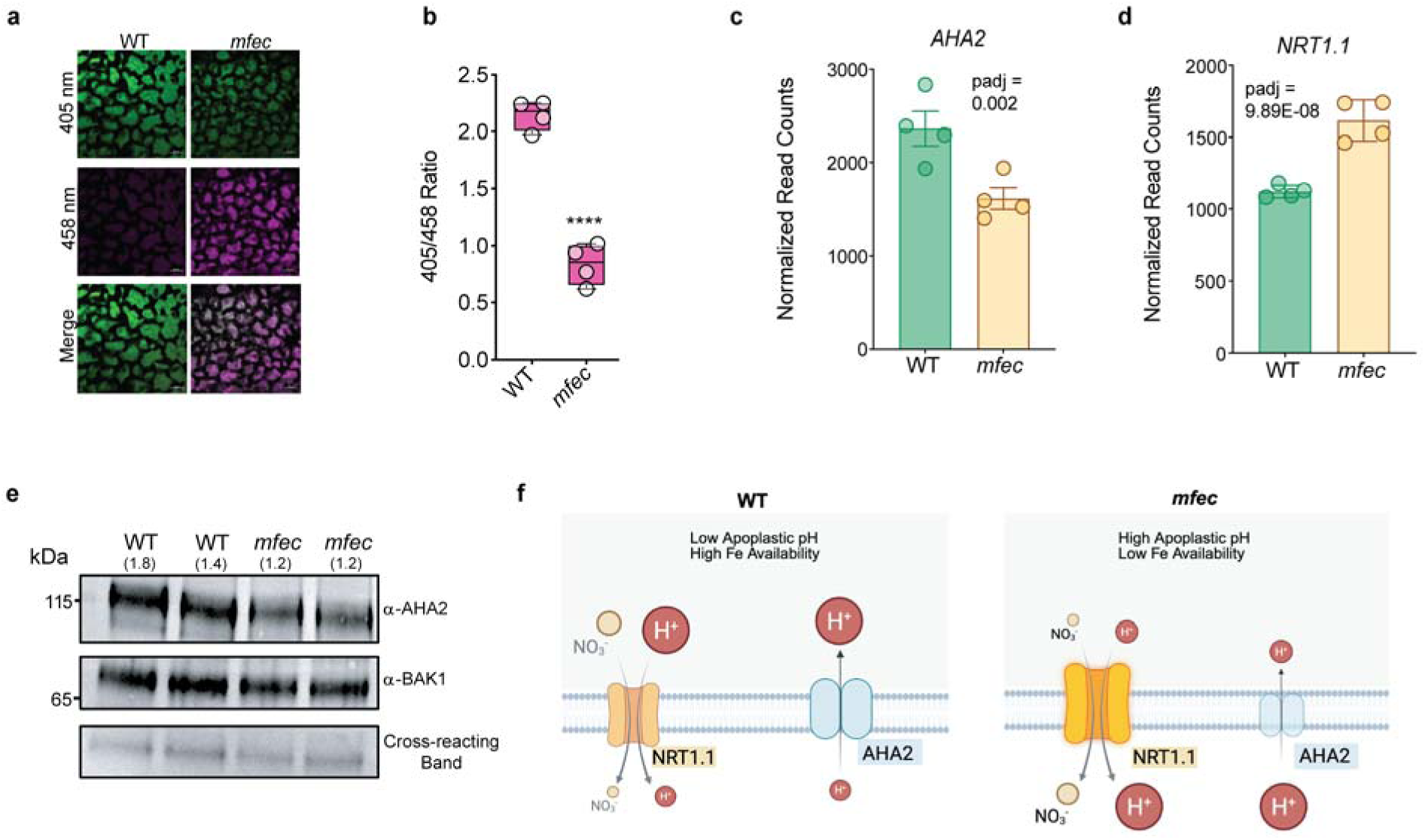
*mfec* mutants exhibit altered apoplastic pH regulation through dysregulated AHA2 level. **a**, Representative HPTS ratiometric images of apoplastic pH in WT and *mfec* leaves. **b**, Quantification of apoplastic pH from HPTS fluorescence ratio (405/458 nm excitation). **** indicates p < 0.0001 determined by Student’s t-test. **c,d** Normalized RNA sequencing read counts of (**c**) *AHA2* and (**d**) *NRT1.1* transcripts in WT and *mfec* leaves from 4.5-week-old plants grown in potting soil. Each dot represents one biological replicate (n = 4); adjusted p-value (padj) from DESeq2 Wald test with Benjamini-Hochberg correction for multiple testing. Bars represent mean ± SEM. **e,** Western blot of AHA2 protein abundance in WT and *mfec* leaves (two biological replicates per genotype shown), with BAK1 as a membrane loading control, and a non-specific cross-reacting band as an additional loading control. Molecular weight markers (kDa) shown at left. Numbers in parentheses above each lane indicate AHA2 band intensity normalized to BAK1. Source data for unprocessed blot is available in Supplemental Fig. 1c. **f**, Schematic model of apoplastic ion transport in WT and *mfec* leaves. In WT leaves, low apoplastic pH and high Fe availability are associated with NRT1.1-mediated NO_3_ /H symport and AHA2-mediated H efflux from the apoplast. In *mfec*, high apoplastic pH and low Fe availability are correlated with increased *NRT1.1* abundance and reduced *AHA2* level.

### Restoring iron re-establishes microbiota homeostasis

If reduced iron bioavailability drives microbiota dysbiosis in *mfec* leaves, we reasoned that restoring soluble iron might reverse both microbial over-proliferation and bacterial community imbalance. We therefore tested whether supplementation with bioavailable iron is sufficient to suppress the dysbiotic phenotypes. Consistent with our previous observations, *mfec* leaves harbored substantially greater bacterial loads than WT plants following inoculation with the leaf synthetic community (SynCom^Col-0^) (Fig. 4a,b). Remarkably, supplementation with 5 μM FeSO_4_, which falls within the physiological range measured in the leaf apoplastic fluid of iron-sufficient plants [28], suppressed endophytic bacterial abundance in *mfec* leaves to levels comparable to those in WT leaves, whereas iron treatment had little effect on endophytic microbiota abundance in WT plants (Fig. 4b). Thus, increasing iron bioavailability is sufficient to suppress the excessive bacterial proliferation characteristic of the dysbiotic *mfec* mutant. The restoration of the microbiota level was accompanied by recovery of plant health. Whereas untreated *mfec* plants developed the characteristic chlorosis and necrosis, iron supplementation largely abolished these symptoms, resulting in plants that closely resembled WT leaves (Fig. 4a). These observations confirmed that restoring iron availability simultaneously rescues regulation of endophytic leaf microbiota level and plant health.

In addition to microbiota load, we also analyzed the 16S rRNA profiles under standard and iron supplemented conditions (Fig. 4c). In these experiments, the endophytic compartment of WT leaves was dominated by *Pandoraea* Amplicon Sequence Variants (ASVs) and the community composition remained similar regardless of Fe^2+^ treatment (Fig. 4c). In contrast, *mfec* leaves showed increased relative abundance of *Achromobacter*, *Comamonas* and *Stenotrophomonas* ASVs and reduced relative abundance of *Pandoraea* ASVs (Extended Data Fig. 3a-d). Strikingly, Fe^2+^ supplementation restored the *mfec* microbiota composition toward that of WT, as observed by the increased *Pandorea* and *Paenibacillus* ASVs and reduced *Achromobacter*, *Comamonas* and *Stenotrophomonas* ASVs even though *Paenibacillus* showed a low abundance in both WT and *mfec* microbiota profiles in these experiments

### Iron availability controls microbial infighting

Previous work showed that dysbiosis in the *mfec* mutant is characterized by expansion of Gammaproteobacteria, including *Stenotrophomonas*, *Achromobacter* and *Comamonas* and depletion of Firmicutes such as *Paenibacillus* due to widespread Proteobacteria-Firmicute infighting [5]. The reduced iron bioavailability in *mfec* leaves prompted us to ask whether iron limitation directly influences interactions among members of the leaf microbiota. To investigate this hypothesis, we first used FeGenie [23] to annotate iron-related genes across all 48 members of SynCom^Col-0^ including C46 (Fig. 4d). This analysis revealed that most iron acquisition and regulatory functions, including siderophore synthesis, siderophore transport, and iron transport, were broadly distributed across all four phyla represented in SynCom^Col-0^ and C46. However, Firmicutes lacked the TonB-dependent iron transport system (annotated by FeGenie as ’iron acquisition-siderophore transport potential) present in the three Gram-negative phyla, consistent with their lack of an outer membrane [24]. This pattern was corroborated by antiSMASH analysis [25] of predicted siderophore biosynthetic gene clusters, which identified the majority of siderophore-producing potential within Gammaproteobacteria (21 of 29 predicted siderophore biosynthetic gene clusters (BGCs)), with substantially fewer in Firmicutes (6) and Alphaproteobacteria (2), and none detected in Bacteroidetes (Fig. 4e, Supplementary Table S6, S7). This enrichment was not simply a consequence of Gammaproteobacteria’s larger representation in the community, as siderophore BGCs per strain remained higher in Gammaproteobacteria than in Firmicutes or Alphaproteobacteria after normalizing by strain number within each phylum

Because TonB-dependent receptors enable Gram-negative bacteria to import siderophores produced by other taxa [26], Firmicutes lacking this system may be specifically limited in their ability to access iron sequestered by siderophore-producing Gammaproteobacteria, even though they retain other iron acquisition and transport capabilities. To investigate this possibility, we performed *in vitro* binary interactions between the Proteobacterium *S. maltophilia* C46 and the Firmicute *Paenibacillus chondroitinus* C3, a strain previously shown to be inhibited by C46 [5]. As an inhibition-negative control, we included a closely related *P. chondroitinus* strain (C17-2), which was previously shown not to be inhibited by C46 [5]. We reproduced the observation that C46 inhibits C3 but not C17-2 on R2A medium (Fig. 4f, Extended Data Fig. 4 a,b). Given this difference in inhibition between two closely related strains, we performed comparative genome analysis between C3 and C17-2 to look for genomic features that could explain the differential susceptibility to C46. We hypothesized that despite their ∼99% genome similarity, C17-2 might encode genes enabling it to coexist with C46 without being outcompeted. Using RAST subsystem annotation, we found that C17-2 uniquely encodes a heme-iron acquisition and detoxification module, comprising the heme efflux transporter HrtAB, an iron regulated surface determinant (Isd)-like heme transport system (*IsdC*, *IsdE*, *IsdF*, and an associated ATP-binding component), and two sortases (a housekeeping sortase A and an NPQTN-specific sortase B typically associated with Isd-type systems) (Supplementary Table S5). These genes are entirely absent from the C3 genome. To test whether this difference in iron acquisition capacity underlies the antagonism of C3, we supplemented R2A containing 2,2^’^-bipyridyl (DIP) with iron. Iron supplementation alleviated inhibition of C3 by C46, demonstrating that C46 inhibits C3 in an iron-dependent manner (Fig. 4e, Extended Data Fig. 4 a,b). As expected, C46 did not inhibit C17-2 on R2A plate with or without iron supplementation (Extended Data Fig. 4b).

We extended the *in vitro* iron-dependent binary competition assay to all 31 Proteobacteria strains within SynCom^Col-0^ and found that 26 of the 31 Proteobacteria strains inhibited C3 and iron supplementation alleviated inhibition for 23 of these 26 inhibitory strains (Extended Data Fig. 4a). This pattern was consistent across *Stenotrophomonas*, *Comamonas*, and *Variovorax,* all of which showed larger zones of clearing against C3 which decreased substantially upon iron supplementation. Chrome Azurol S (CAS) assays [27] showed that siderophore production in these strains including C46 was induced under iron-depleted conditions (Extended Data Fig. 4a,c). In contrast, *Achromobacter* strains inhibited C3 and produced strong siderophores but this inhibition was not alleviated by iron supplementation (Extended Data Fig. 4a). This suggests that the C3-inhibitory activity of *Achromobacter*, which is strongest under low iron availability, may be mediated via an iron competition-independent mechanism.

Finally, to determine if the iron-dependent Proteobacteria-Firmicute inhibition occurs *in planta*, we infiltrated C46 and C3 together with 5 µM soluble iron (FeSO_4_) into leaves of 4-week-old Arabidopsis WT plants. Consistent with our *in vitro* findings, co-infiltration with C46 reduced C3 levels, relative to C3 infiltrated alone (Fig. 4g). Iron supplementation restored C3 levels to those observed with C3 alone, indicating that C46 inhibits C3 in an iron-dependent manner in planta (Fig. 4g). On the other hand, the population of C46 was unaffected by the presence of C3 and remained stable whether C46 was infiltrated alone or with C3, and regardless of iron supplementation (Fig. 4h).

**Figure 4.**
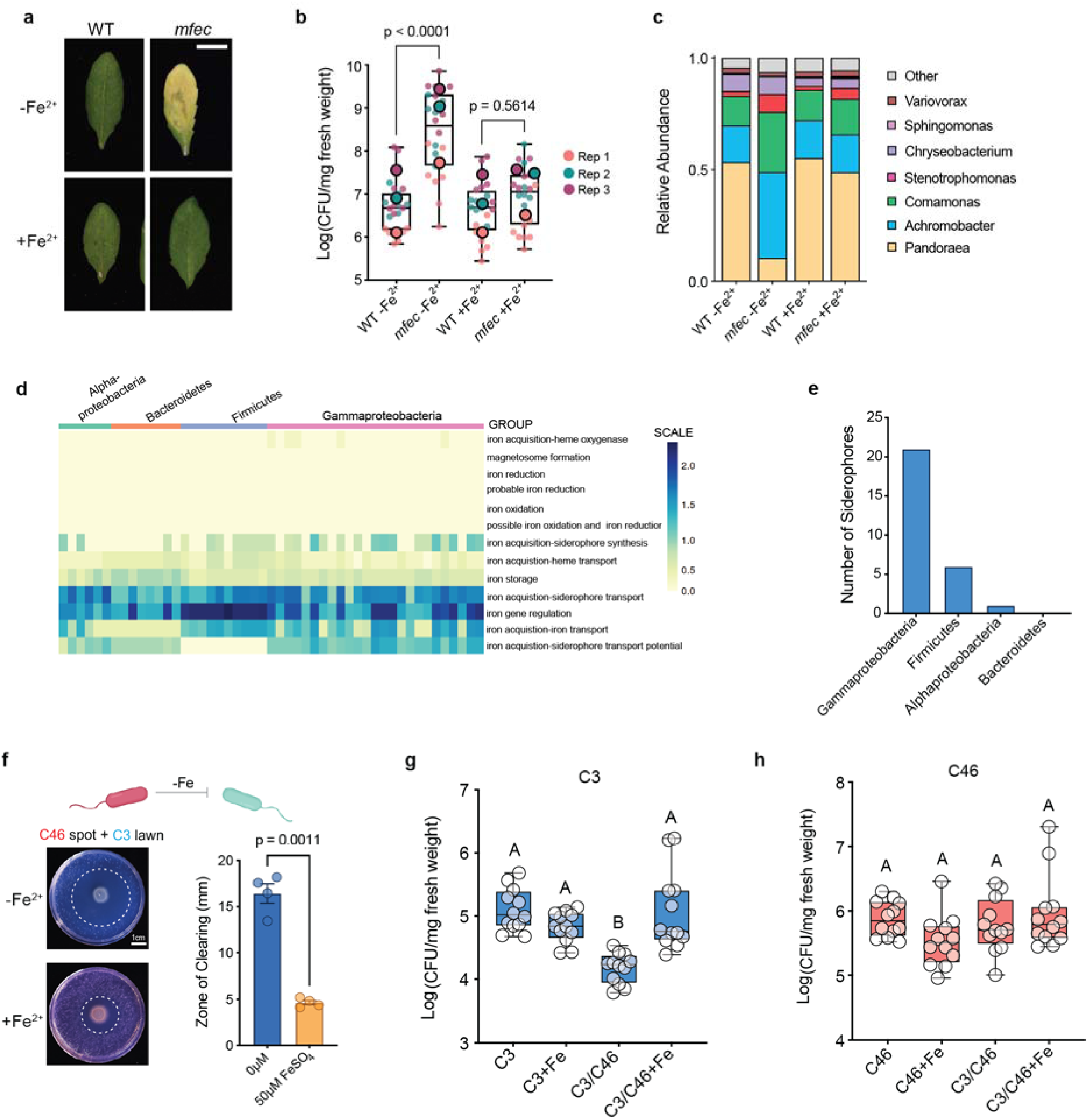
Iron supplementation restores microbiota homeostasis and alleviates dysbiotic symptoms in *mfec* leaves. **a**, Representative images of WT and *mfec* leaves infiltrated with SynCom^Col-0^ with or without supplemented Fe^2+^ (5 µM FeSO_4_), after 4 days of >95% relative humidity treatment. **b**, Endophytic microbiota counts in WT and *mfec* leaves, +/-5 µM Fe^2+^ supplementation, following SynCom^Col-0^ infiltration as in (**a**). Small dots represent biological replicates within an experiment and large, outlined dots represent mean value per experiment (n = 3 independent experiments colored by replicate and eight biological replicates per experiment). Statistical significance determined by one-way ANOVA (p < 0.0001) with Tukey’s multiple comparisons test; p-values shown for WT vs. *mfec*. **c**, Relative abundance by genus from 16S rRNA amplicon profiling from WT and *mfec* leaves +/-5 µM Fe^2+^ supplementation. **d**, Heatmap of iron acquisition and regulatory gene functions annotated by FeGenie across members of SynCom^Col-0^, grouped by phylum, with rows (functional categories) hierarchically clustered. Color scale indicates the percentage of each genome’s protein-coding genes (ORFs) assigned to each iron-related functional category, per FeGenie’s standard normalization (gene count for category/total ORF count). **e**, Number of predicted siderophore biosynthetic gene clusters (BGCs) per phylum, identified by antiSMASH analysis of SynCom^Col-0^ genomes. **f**, *In vitro* binary interaction assay between *S. maltophilia* C46 (spot) and *P. chondroitinus* C3 (lawn) on R2A medium containing 2,2′-bipyridine with or without Fe^2+^. Representative images (left) show zones of clearing around C46. Quantification of zone of clearing without (0 µM) vs with (50 µM) FeSO_4_ (right). Each dot represents one biological replicate; bars show mean ± SEM; p-value from a two-tailed t-test. **g,h**, *In planta* bacterial titers of (**g**) C3 and (**h**) C46 in WT leaves infiltrated with each strain alone (control), with Fe (control), in combination with the other strain (C3/C46), or in combination with the other strain plus 5 µM Fe^2+^. Data are from three independent experiments, each with four biological replicates (n = 12 per group). Statistical significance in (**g,h**) was determined by one-way ANOVA with Tukey’s multiple comparisons test; different letters indicate statistically significant groups.

**Extended Data Fig. 3.**
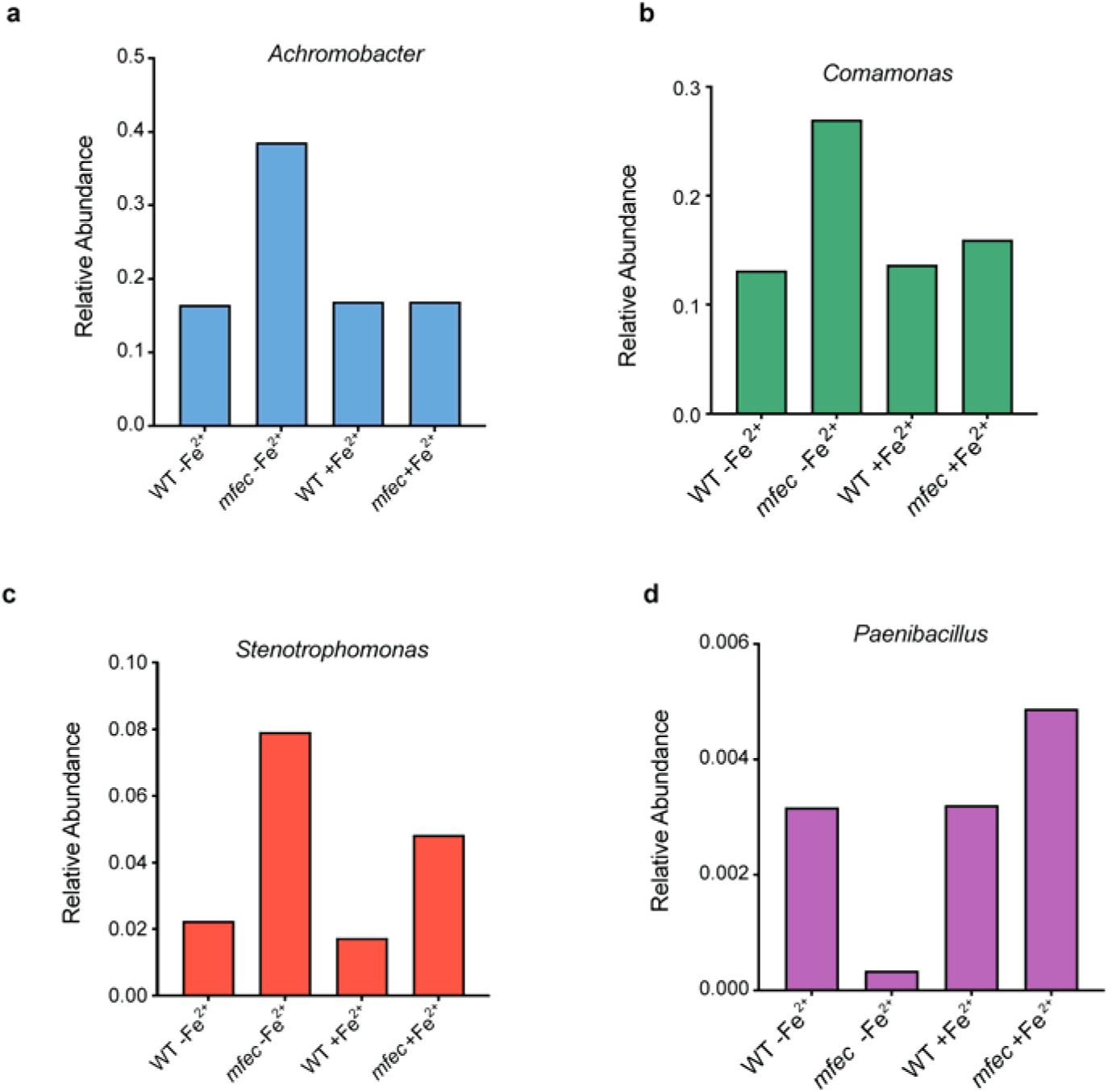
Iron supplementation shifts individual taxon relative abundance in *mfec* leaves toward that of WT. Relative abundance of (**a**) *Achromobacter*, (**b**) *Comamonas*, (**c**) *Stenotrophomonas*, and (**d**) *Paenibacillus* amplicon sequence variants (ASVs) from 16S rRNA profiling of WT and *mfec* leaves, with or without 5 µM Fe^2+^ supplementation, as in Fig. 4c. Bars represent relative abundance pooled across biological replicates for each condition.

**Extended Data Fig. 4.**
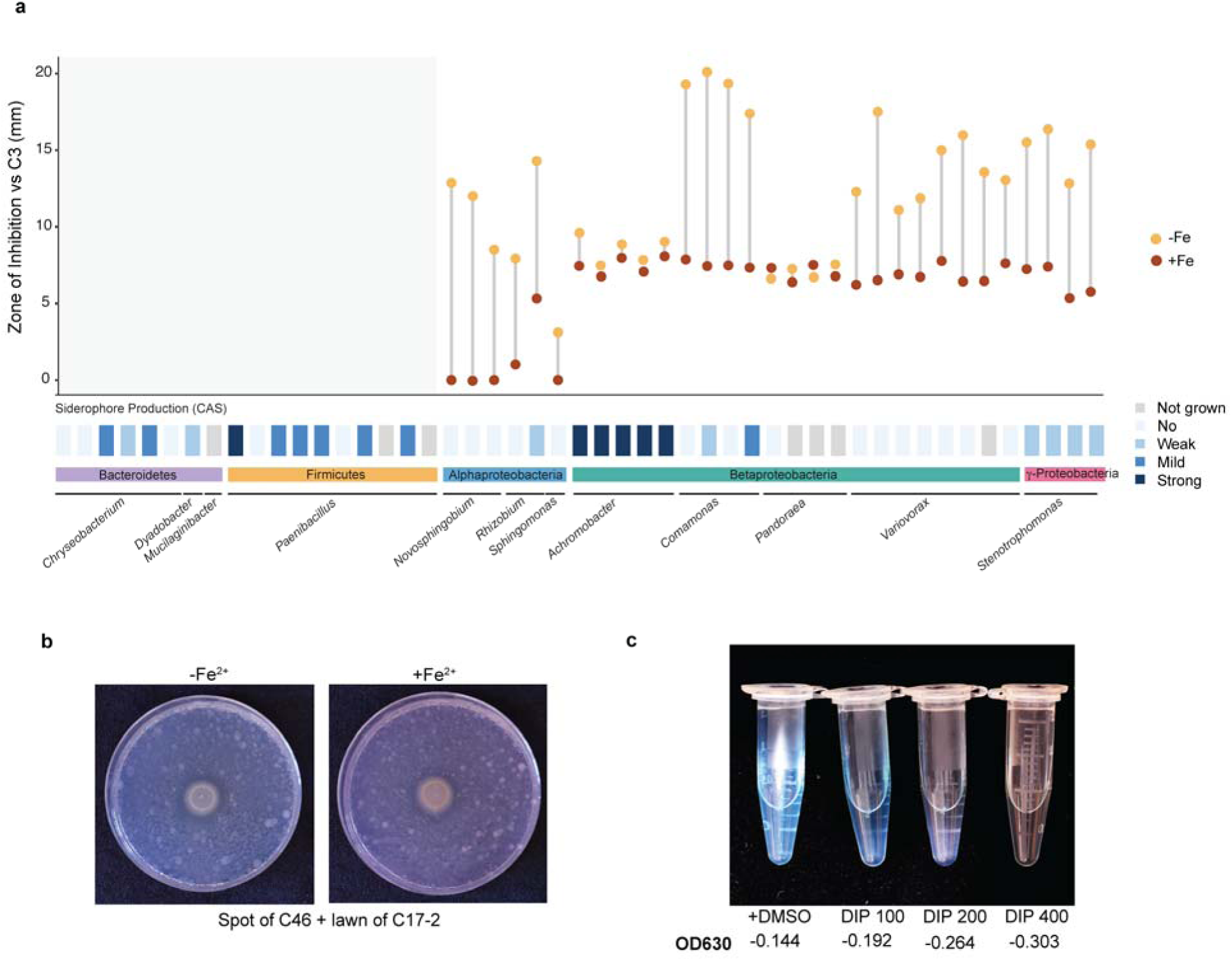
Iron-dependent inhibition of *P. chondroitinus* (C3) among Proteobacteria in SynCom^Col-0^ and their siderophore production. **a**, (Top) Zone of inhibition (mm) against C3 in *in vitro* binary competition assays for 31 Proteobacteria strains within SynCom^Col-0^, grown on R2A medium containing 2,2′-bipyridine (DIP) with (+Fe, dark orange) or without (-Fe, light orange) 50 µM FeSO_4_ supplementation grouped by phylum and genus. Binary interaction assays were performed only for the Proteobacteria subset. (Bottom), Chrome Azurol S (CAS) siderophore production score for each strain (not grown, No, Weak, mild, strong is used to indicate strength of inhibition). **b,** Representative images of C46 spotted onto a lawn of the non-inhibited control strain *P. chondroitinus* C17-2 on R2A + DIP, with (+Fe^2+^) or without (-Fe^2+^) 50 µM FeSO_4_ supplementation, as in Fig. 4f. **c,** Chrome Azurol S (CAS) liquid assay of siderophore production under increasing iron chelation (+DMSO control, and DIP at 100, 200, and 400 µM); OD_630_ values shown below each tube, with decreasing OD_630_ indicating greater dye decolorization and increasing siderophore activity.

### Iron-mediated dysbiosis extends to pathogen infection

The *mfec* mutant was originally constructed to genetically mimic the disruption of PRR signaling and MIN7-dependent vesicle trafficking caused by *Pseudomonas syringae* type III effectors [17]. We next tested the hypothesis that the iron-dependent dysbiosis observed in *mfec* leaves represents a natural feature of bacterial infection rather than a mutant-specific phenomenon. We first investigated whether *Pst* DC3000 directly competes with resident leaf microbiota for iron by conducting *in vitro* binary interaction studies. We found that, like other Gammaproteobacteria leaf microbiota, *Pst* DC3000 inhibited *P. chondroitinus* (C3), and exogenous iron supplementation alleviated inhibition of C3 by *Pst* DC3000 (Fig. 5a). Furthermore, a *Pst* DC3000 triple mutant deficient in siderophore-mediated iron acquisition, lacking citrate uptake (*fecA*), pyoverdine (*pvdI*), and yersiniabactin (*pvhA*) [29], had substantially reduced inhibitory activity against C3 compared to WT *Pst* DC3000. We next expanded the *Pst* DC3000-leaf microbiota binary interaction to include all 272 SynCom strains from two independent SynComs isolated from *Arabidopsis* leaves (He Lab and Vorholt Collection). Remarkably, approximately 40% of leaf microbiota strains were inhibited by *Pst* DC3000 in an iron-dependent manner (Fig. 5b, Extended Data Fig. 5a, Supplementary Table S8), indicating that iron competition is a key mechanism by which *Pst* DC3000 antagonizes resident leaf microbiota during pathogenesis.

Next, we examined whether *Pst* DC3000 infection disrupt leaf microbiota *in planta* and, if yes, if siderophore plays a role in microbiota disruption *in planta*. We noted that *Arabidopsis* Col-0 WT plants infiltrated with the *Pst* DC3000 Δ*fecA*Δ*pvdI*Δ*pvhA* mutant showed less disease susceptibility compared to when infected with WT *Pst* DC3000, and the Δ*fecA*Δ*pvdI*Δ*pvhA* mutant grew less *in planta* compared to WT *Pst* DC3000, indicating that siderophore-mediated iron acquisition contributes to *Pst* DC3000 virulence (Fig. 5c,d). Notably, the *Pst* DC3000 Δ*fecA*Δ*pvdI*Δ*pvhA* grew similarly to WT *Pst* DC3000 *in vitro*, indicating that its reduced inhibitory activity was not attributable to a general growth defect (Extended Data Fig. 5b). As expected, plants infected with the T3SS-deficient *hrcC* mutant showed no disease symptoms and correspondingly low pathogen counts. Using 16S rRNA profiling, we found that WT *Pst* DC3000 infection drove Gammaproteobacteria enrichment and Bacilli depletion, a community shift strikingly reminiscent of that observed in *mfec* plants [5]. In contrast, the *fecA pvdI pvhA* mutant produced an intermediate composition between WT and mock (Fig. 5e). Together, these findings suggest that *P. syringae* promote microbiota dysbiosis through two complementary mechanisms. First, T3SS effector-mediated disruption of PRR signaling and MIN7-dependent trafficking, as mimicked by the *mfec* mutant, creates an iron-limited environment that favors dysbiosis. Second, the pathogen additionally exploits siderophore-mediated competition that further disrupts resident commensal bacteria.

**Figure 5.**
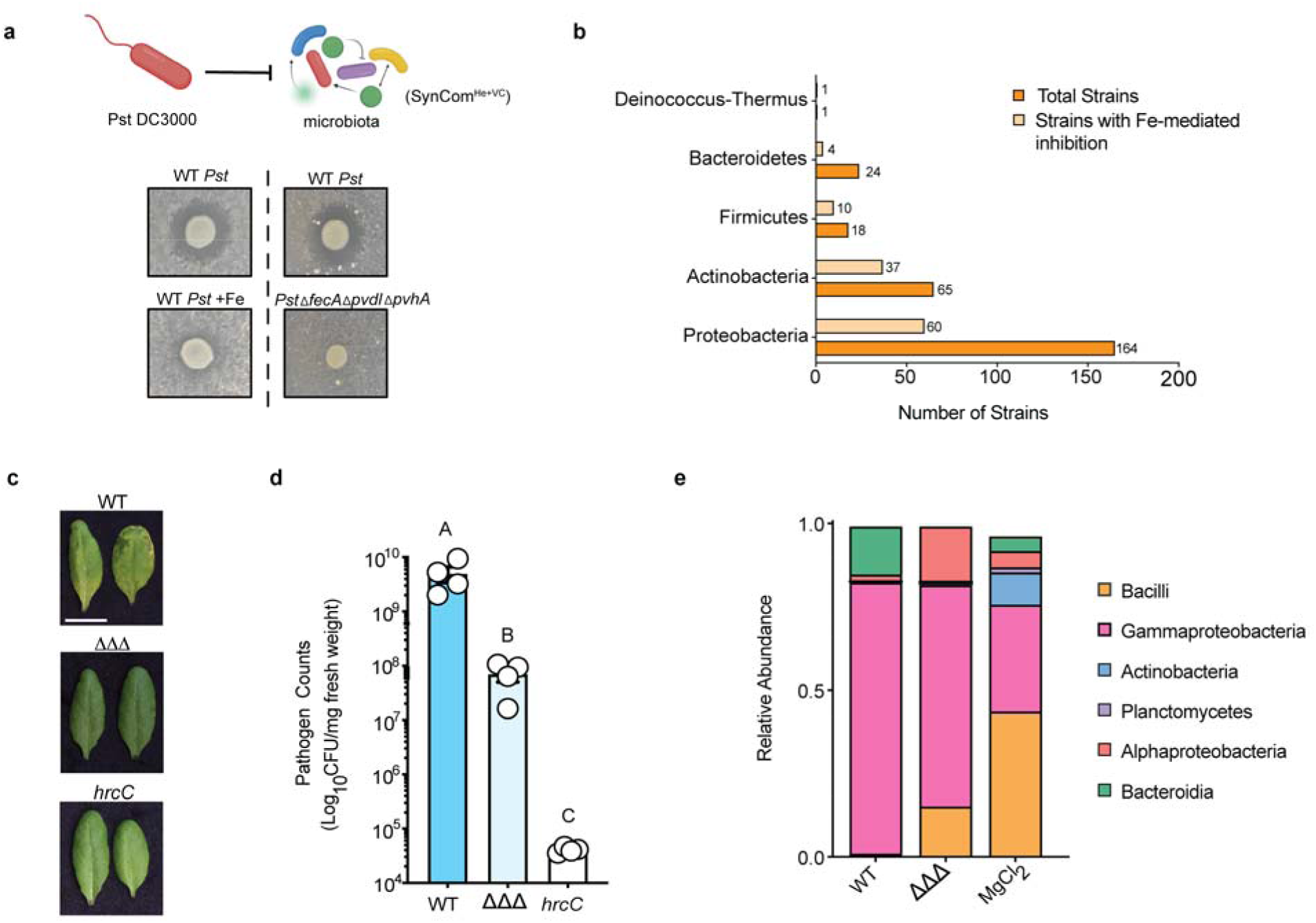
*Pst* DC3000 iron acquisition machinery contributes to leaf microbiota dysbiosis and virulence. **a**, (Top) Schematic illustrating *Pst* DC3000-mediated inhibition of microbiota. (Bottom) Representative zone-of-inhibition plate assays: WT *Pst* DC3000 spotted against a *P. chondroitinus* C3 lawn, with (+Fe) or without (-Fe) iron supplementation (left), and WT *Pst* DC3000 versus the iron-acquisition-deficient *Pst* DC3000 Δ*fecA* Δ*pvdI* Δ*pvhA* triple mutant spotted against a C3 lawn (right). **b**, Number of microbiota strains, grouped by phylum, showing iron-mediated inhibition by *Pst* DC3000. Bright orange bars indicate total strains tested per phylum; light bars indicate the subset showing Fe-mediated inhibition (i.e., inhibition relieved by iron supplementation). Numbers above each bar indicate strain counts. **c**, Representative leaf images from WT, Δ*fecA* Δ*pvdI* Δ*pvhA* (ΔΔΔ), and *hrcC Pst* DC3000 infections, showing disease symptoms. **d**, Endophytic pathogen counts in leaves infected with *Pst* DC3000 WT, ΔΔΔ, or *hrcC*. Statistical significance determined by One-way ANOVA with Tukey’s multiple comparisons test; different letters indicate statistically significant groups. Experiment performed three independent times with at least three biological replicates per treatment. e, 16S rRNA relative abundance of leaf-associated bacterial communities following infiltration with WT *Pst* DC3000, Δ*fecA* Δ*pvdI* Δ*pvhA* or MgCl_2_ (mock infiltration control).

**Extended Data Fig. 5.**
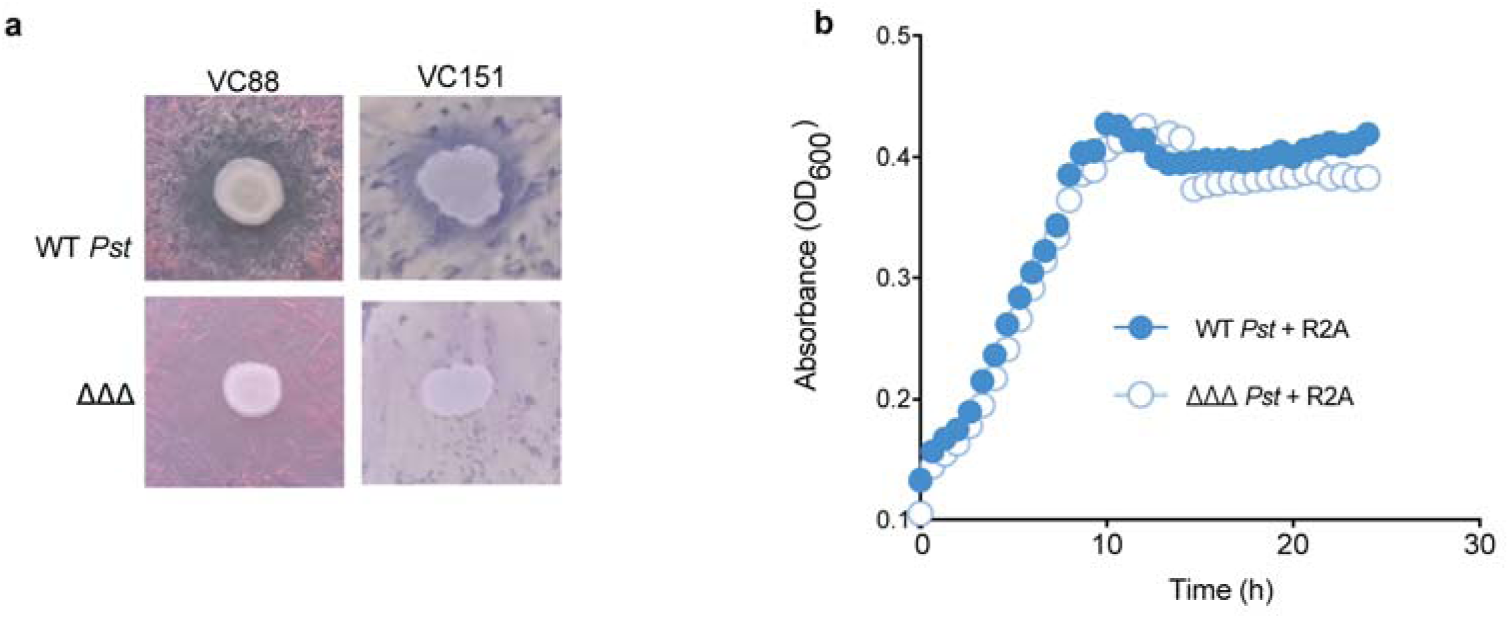
Siderophore-mediated iron acquisition promotes pathogen antagonism of leaf microbiota. **a**, Binary interactions showing inhibition of three microbiota strains from the Vorholt Collection (VC88 and VC151) by WT *Pst* DC3000 and the siderophore-deficient Δ*fecA* Δ*pvdI* Δ*pvhA* mutant. **b,** Growth curves of WT *Pst* DC3000 (filled circles) and the *Pst* DC3000 Δ*fecA* Δ*pvdI* Δ*pvhA* mutant (open circles) in R2A medium over 24 h, measured at OD_600._

## DISCUSSION

A fundamental question in host-microbiota biology is how hosts selectively assemble a healthy microbial community while limiting the proliferation of opportunistic or pathogenic microorganisms. Although previous studies demonstrated that plant immune signaling, vesicle trafficking and other cellular processes are required to maintain phyllosphere microbiota homeostasis [4–6], the metabolic mechanisms through which these host genes/pathways shape phyllosphere microbial communities and prevent dysbiosis remained largely unknown. Following the KEGG-derived clues from the transcriptomic analysis of a prominent dysbiosis-inducing endophytic bacterial strain (*S. maltophilia* C46), we found that iron bioavailability plays a critical role in shaping healthy vs. dysbiotic microbiota assembly in the Arabidopsis leaf endosphere. Importantly, the iron bioavailability-based mechanism appears to be integral to infection-associated dysbiosis induced by a common foliar pathogen, *P. syringae*. These results uncovered a prominent role for iron as a determinant in mediating a switch between healthy and dysbiotic endophytic leaf microbiota.

Our findings suggest that endophytic leaf microbiota homeostasis depends on microbial access to limiting iron resources rather than the absolute abundance of iron within leaf tissues. We did not find a drastic difference in total iron levels between WT and the *mfec* mutant. However, the transcriptomes of *Stenotrophomonas* strains clearly indicated an iron starvation endophytic environment. Iron supplementation is sufficient to recover a healthy microbiota and abates dysbiotic symptoms, demonstrating that iron accessibility is itself a host-regulated determinant of endophytic leaf microbiota. This distinction between nutrient abundance and nutrient availability has broad implications for understanding host-associated microbial communities, where microbes may respond primarily to the local chemical environment rather than bulk nutrient content.

The elevated apoplastic pH in *mfec* leaves is consistent with low endophytic iron bioavailability sensed by *Stenotrophomonas* strains. However, future research should investigate if the elevated apoplastic pH could impact additional host-controlled parameters, including the availability of other micronutrients, carbon sources, redox status, or water potential, which could additionally contribute to microbiota homeostasis in the phyllosphere. We found that PRR immune signaling and MIN7-dependent vesicle trafficking affect the level expression of the H⁺-ATPase AHA2 (and likely other immune-associated plasma membrane cargoes) as part of a plant genetic network to maintain endophytic phyllosphere microbiota homeostasis. This finding suggests that immune signaling, vesicle trafficking, pH homeostasis, and iron availability may not be independent processes but interconnected components of a regulatory network that establishes a stable endophytic niche for a healthy microbiota community while restricting dysbiosis.

In this study, we found that a common foliar pathogen, *P. syringae*, which targets PRR immune signaling and MIN7-dependent vesicle traffic using its T3SS effectors [17], appears to simultaneously perturbs host pathways that reduce iron bioavailability and competes aggressively for the remaining accessible iron through siderophore-mediated acquisition. These complementary iron-depleting activities suggest that successful pathogens, like *P. syringae*, may have evolved to manipulate not only host immune signaling but also direct microbe-microbe interactions that governs microbial competition. It would be interesting in the future to investigate if pathogen-induced dysbiosis represent simply collateral damages of infection or influence, positively or negatively, pathogenesis and/or pathogen persistence in natural cycles.

Broadly, our work supports a conceptual framework that regulation of iron bioavailability may represents a general strategy through which plants shape a healthy leaf microbiota, with pathogens evolving counterstrategies that exploit the same molecular principles to promote colonization and disease. Notably, iron availability is also a key determinant of human host--microbiota interactions [30, 31]. Thus, the role of iron in maintaining microbiota homeostasis and dysbiosis prevention may be universal across the kingdoms of life.

## Supporting information

Supplementary Western blot images

Suppmentary Table 1-8

## MATERIALS AND METHODS

### Plant Materials and Growth Conditions

Arabidopsis thaliana seeds were sterilized with 10% bleach and washed 2-3 times with autoclaved milliQ water, then stratified at 4℃ for 2 days. Stratified seeds were grown in 3.5^’’^ pots with Arabidopsis mix potting soil (1:1:1 SureMix (Michigan Grower Products): vermiculite: perlite) in air circulating growth chambers at ∼60% relative humidity, 20-22°C, 12h:12h light:dark photoperiod, and a light intensity of 100 μmol m^-2^ s^-1^ (standard conditions). Seeds were allowed to germinate for one week at 95% humidity (flats covered with a dome), with ambient humidity thereafter for 4-4.5 weeks. Plants were fertilized with 20-20-20 fertilizer during the first week and watered with water alone twice per week thereafter. Four to four and half week-old plants were used for all experiments unless otherwise stated.

The wild-type accession Col-0, *min7 fls2 efr cerk1* (*mfec*) was obtained and described in previous studies [1, 2]. All images were captured using a Nikon AF-S DX Micro-NIKKOR 40mm f/2.8G camera.

### Preparation of synthetic community

Isolates of SynCom^Col-0^ and C46 were streaked from glycerol stocks on R2A (Sigma, 17209) plates and allowed to grow at room temperature for two days. Isolates were scraped from plates and resuspended in 500 µl 10 mM MgCl_2_. Cultures were normalized to the strain with the lowest OD_600_. The cultures were combined and normalized to a final OD_600_ of 0.04 which corresponds to ∼10^7^ CFU/mL.

### Bacterial *in planta* transcriptomics

Leaves of four-and-a-half-week-old WT and *mfec* Arabidopsis plants were syringe-infiltrated with either *Stenotrophomonas maltophilia* (C46) alone or SynCom^Col-0^. Bacterial cultures were normalized to OD_600_ of 0.9. Plants were allowed to dry after infiltration for 30 mins and placed in a growth chamber at standard conditions but with >95% humidity for 6 h. Leaves were collected in 50 mL conical tubes containing 8-10 steel beads, flash-frozen in liquid nitrogen and stored at -80°C until bacterial isolation and RNA extraction. The experiment included four conditions - C46 alone in WT, C46 alone in *mfec*, SynCom^Col-0^ with C46 in WT, and SynCom^Col-0^ with C46 in *mfec*. Four leaves per plant were infiltrated and each condition contained 4 biological replicates giving 16 samples total.

#### Bacterial Isolation

Bacteria were isolated based on a protocol developed in Nobori *et al.* 2018 but with modifications [3]. Frozen leaves in tubes were initially shaken vigorously by hand to crush the leaves into small pieces and a 1 mL pipette tip was used to aid further crushing. Crushed leaves were ground to powder using a mortar and pestle under liquid nitrogen. Thirty milliliters of ice cold ice-cold bacterial isolation buffer Nuclease-Free Water, 25 mM TCEP (tris(2-carboxyethyl)phosphine) pH 4.5 adjusted with 10 N NaOH, 9.5% ethanol,0.5% phenol solution in a fume hood. Samples were vortexed immediately for 10 s, followed by vigorous shaking by hand for 10 s, and then incubated for 20 h with shaking on a Roller Mixer SRT3 at 4°C. The resulting mixture was filtered through a 40 µm cell strainer into a new 50 mL tube on ice, using 2-3 strainers per sample as needed. The flow-through was centrifuged for 20 mins at 3,200 × *g* at 4°C. The supernatant was removed using a serological pipette and ∼500 µL of liquid was left behind to avoid sample loss. Nine hundred microliters of ice-cold ethanol were added, and the pellet was fully resuspended by vortexing. The sample was then transferred to a new 1.5 mL tube on ice. Samples were centrifuged for 20 mins at 2,300 × *g* at 4°C to yield a two-layered pellet (white on top, green underneath). Using a 20 µl pipette set to 18 µL, the top (white) layer was carefully resuspended; once the buffer became cloudy, the suspension was transferred to a new 1.5 mL tube. Bacterial cells were harvested by centrifugation for 2 mins at 10,000 × *g*, and the supernatant was discarded.

#### RNA Extraction

RNA was extracted from bacterial pellets using the Direct-zol RNA Miniprep Kit (Zymo Research). Pellets were resuspended in 100 µL lysozyme solution (50 mg/mL) per sample, vortexed at maximum speed for 30s and incubated at 30°C for 10 min. Three hundred microliters of TRI Reagent® (Trizol; 3 volumes per 1 volume of cell suspension) was then added and samples were vortexed at maximum speed for 5 mins. An equal volume (400 µL) of 95-100% ethanol was added to each Trizol-lysed sample and mixed thoroughly. RNA was purified using the Direct-zol RNA Miniprep Kit (Zymo Research) according to the manufacturer’s instructions, with an in-column DNase I treatment using the RNase-Free DNase Set (Qiagen) to remove contaminating genomic DNA. RNA was eluted by adding 50 µL DNase/RNase-free water directly to the column matrix.

### Library Preparation and Sequencing

Ribosomal RNA was depleted from total RNA using the QIAseq FastSelect −rRNA Plant Kit and the FastSelect − 5S/16S/23S Kit (Qiagen) in combination, per manufacturer’s instructions. Sequencing libraries were prepared using the Illumina TruSeq Total RNA Library Preparation Kit with IDT for Illumina Unique Dual Index adapters, following the manufacturer’s recommendations. Completed libraries were quality-checked and quantified using the Qubit dsDNA HS Assay (Thermo Fisher Scientific) and the Agilent 4200 TapeStation HS DNA1000 assay, then pooled in equimolar amounts and the pool quantified using the KAPA Biosystems Illumina Library Quantification qPCR Kit. The pooled library was sequenced on a single lane of an Illumina HiSeq 4000 in 2 × 150 bp paired-end format using HiSeq 4000 SBS reagents. Base calling was performed with Illumina Real-Time Analysis (RTA) v2.7.7, and reads were demultiplexed and converted to fastq format using Illumina bcl2fastq v2.19.1. Sequencing was performed at the Research Technology Support Facility Genomics Core, Michigan State University. Raw sequencing data are available under NCBI SRA SRA project ID PRJNA864025

### Bacteria RNA-sequencing and data analysis

Raw RNA-seq data were filtered by removing low-quality reads and adapters using sickle (v1.33; https://github.com/najoshi/sickle). The high-quality read was aligned to the reference genome of *S. maltophilia* (C46) and other 47 strains (NCBI Bio project ID PRJNA864025) using HISAT2 version 2.2.1 [4] and SAMtools version 1.2 [5]. The gene prediction and annotation of C46 genome was performed using Prokka pipeline version 1.12 [6] and Kyoto Encyclopedia of Genes and Genomes database (KEGG) [7]. Each gene was quantified using HTSeq-count version 0.11.2 [8]. The differentially expressed genes (DEGs) were determined using DESeq2 version 1.30.1 [9] and packages in R version 4.2. The enrichment analysis of KEGG functional pathways of DEGs was performed using Fisher’s exact test based on an adjusted *P* ≤ 0.05. The PCoA analysis was performed based on Bray-cutis distance using vegan package version 2.6-6.1 [10] under R version 4.2.

### Plant RNA-sequencing

Leaves from Col-0 WT and *mfec* plants grown as described above were harvested at 2.5 and 4.5 weeks. Leaves were immediately flash-frozen in liquid nitrogen following collection and stored at -80°C until RNA extraction. Total RNA was extracted using RNeasy RNA Purification kit (Qiagen) and DNase treated with RNase-free DNase set (Qiagen) and sent to Research Technology Support Facility Genomics Core, Michigan State University for quality control, library preparation and sequencing.

Sequencing libraries were prepared using the Illumina Stranded mRNA Prep, Ligation kit with IDT for Illumina RNA UD Indexes following the manufacturer’s recommendations except that half-volume reactions were used. Completed libraries were quality-checked and quantified using the Qubit dsDNA HS Assay (Thermo Fisher Scientific) and the Agilent 4200 TapeStation HS DNA1000 assay, then pooled in equimolar amounts and the pool quantified using the Invitrogen Collibri Library Quantification qPCR Kit. The pooled library was sequenced across two lanes of an Illumina NovaSeq SP flow cell in 1 × 100 bp single-end format using a 100-cycle v1.5 reagent kit. Base calling was performed with Illumina Real-Time Analysis (RTA) v3.4.4, and reads were demultiplexed and converted to FASTQ format using Illumina bcl2fastq v2.20.0.

Raw reads were trimmed for adapter content and low-quality bases using Trimmomatic v0.39 [11] in single-end mode, using the TruSeq3-SE adapter set (ILLUMINACLIP:2:30:10) with LEADING:3 and TRAILING:3 quality trimming. Trimmed reads were aligned to the *Arabidopsis thaliana* TAIR10 reference genome using STAR v2.5.2b [12]. Gene-level read counts were quantified using featureCounts (Rsubread v2.14.2) [13], with gene models defined by the TAIR10 GTF annotation, at the meta-feature (gene) level, not counting multimapping or multi-overlapping reads and with a minimum overlap of 1 base. Samples include four biological replicates per genotype for each timepoint. Differentially expressed genes between genotypes (WT, *mfec*) and developmental timepoints (2.5, 4.5 weeks) were identified using DESeq2 [9] in R.

### Microbiota extraction and quantification for iron treatment

Col-0 WT and *mfec* seeds were sown in potting soil as described earlier. Soil was inoculated with 20 mL of a soil slurry containing a soil-derived microbiota prepared from three source soils: agricultural soil collected from Michigan State University, East Lansing, Michigan (MSU22) [2,14], a blueberry farm soil in North Carolina (36° 13′ 07.2″ N, 79° 10′ 45.5″ W) [15] and a Washington soil. Leaves were infiltrated with SynCom^Col-0^ normalized to OD_600_ of 0.04. For iron supplementation experiments, 5µM FeSO_4_ (Sigma-Aldrich, CAS RN: 7782-63-0, Lot #: SLCG0468) was added to the SynCom inoculum. Excess inoculum was blotted away and leaves were allowed to dry before being placed at 95% humidity (domed) treatment for 3-4 days. Leaves were then collected, weighed, surface-sterilized with 5% bleach for 1 min, washed twice in autoclaved milliQ water for 1 min each, and placed in 2 mL impact resistant tubes (Fisher Scientific) containing three 3 mm zirconium oxide beads (GlenMills) and 500 µL 10 mM MgCl_2_. Leaf samples were homogenized at 1500 Hz for 1 min using a tissue lyser (Geno/Grinder, Cole-Parmer HG-600). Homogenates were serially diluted and plated on R2A medium using the dribble plate method and incubated at room temperature for two days. CFU/mg fresh weight was calculated as:

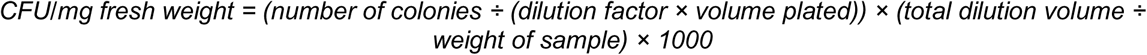

Two leaves were collected per plant and represent one biological replicate. Experiments were performed at least three independent times with eight biological replicates. For 16S rRNA analysis, leaves were collected flash frozen in liquid nitrogen and stored at -80°C until DNA extraction.

### 16S rRNA gene sequencing

We performed 16S rRNA community profiling with Col-0 WT and *mfec* plants treated with and without iron. We also performed 16S rRNA analysis for WT Col-0 plants infected with WT *Pst* DC3000 and *Pst* DC3000 Δ*fecA* Δ*pvdI* Δ*pvhA*. DNA for 16S rRNA extraction and processing was adopted and modified from [2, 15]. Flash frozen tissue samples were ground to a powder and total DNA was extracted with the DNeasy PowerSoil Pro kit (Qiagen) according to the manufacturer’s instructions. PCR was performed in 50 µL reactions using the AccuPrime High-Fidelity Taq DNA Polymerase (Invitrogen) to amplify the V4 region of the bacterial 16S rRNA gene, using the barcoded primer pair 515F-CS1 (5′-ACACTGACGACATGGTTCTACA-GTGYCAGCMGCCGCGGTAA-3′) and 806R-CS2 (5′-TACGGTAGCAGAGACTTGGTCT-GGACTACNVGGGTWTCTAAT-3′) tagged with CS1/CS2 linker sequences for downstream index barcoding by the Michigan State University (MSU) Genomics Core. Peptide nucleic acid (PNA) clamps (PNA Bio) were included to block co-amplification of plant chloroplast and mitochondrial rRNA: pPNA (5′-GGCTCAACCCTGGACAG-3′) and mPNA (5′-GGCAAGTGTTCTTCGGA-3′). PCR products were ran on a 1% agarose gel and the ∼300 bp band of 16S rRNA was excised and extracted using the Zymoclean Gel DNA Recovery kit (Zymo Research). DNA was measured using the PicoGreen dsDNA assay kit (Life Technologies) and all samples were normalized to 2 ng µl^-1^. Samples were submitted to MSU Research Technology Support Facility Research Technology Support Facility Genomics Core. Amplicons were batch-normalized and purified using the Charm Biotech PCR Normalization and Purification Kit, pooled, and the pool was quality-checked and quantified using a combination of the Biotium AccuGreen High Sensitivity dsDNA assay, the Agilent 4200 TapeStation HS DNA1000 assay, and the Invitrogen Collibri Library Quantification qPCR assay. The pooled library was sequenced on a single Element Biosciences AVITI Medium Output flow cell in 2 × 300 bp paired-end format using a Cloudbreak Freestyle 600-cycle reagent kit, with custom sequencing and index primers complementary to the CS1/CS2 oligomers. Base calling was performed using AVITIOS v3.4.2, and reads were demultiplexed and converted to FASTQ format using Element Biosciences bases2fastq v2.2.0.

#### 16S rRNA Analysis

Amplicon libraries targeting the V4 region of the bacterial 16S rRNA gene were generated using the primers 515F (5′-GTGYCAGCMGCCGCGGTAA-3′) and 806R (5′-GGACTACNVGGGTWTCTAAT-3′) [16]. Raw paired-end reads were processed using QIIME2 (version 2025.7) [17]. Primers were removed using cutadapt (DOI:10.14806/ej.17.1.200), discarding any reads in which the primer sequence were not found. The resulting primer-trimmed reads were denoised with DADA2 [18]. Reads were truncated based on per-base quality score profiles, then denoised, dereplicated, merged paired-end reads, checked for chimeras to generate a feature (amplicon sequence variant, ASV) table and corresponding representative sequences with DADA2.

Taxonomy was assigned to representative sequences using a naive Bayes classifier (q2-feature-classifier, classify-sklearn) [19] trained on the SILVA 138.2 SSU NR99 reference database [20]. The reference database was extracted in silico for the 515F/806R amplicon region using RESCRIPt [21]. Reference sequences were quality-filtered using cull-seqs (removing sequences with ≥5 ambiguous bases or homopolymers ≥8 bp) and filter-seqs-length-by-taxon (retaining Archaea ≥900 bp, Bacteria ≥1200 bp, and Eukaryota ≥1400 bp), then dereplicated in “uniq” mode both before and after extracting the primer-specific region.

ASVs classified as chloroplast or mitochondrial in origin were removed from the feature table, and the representative sequence set was filtered to match. Filtered representative sequences were then used to construct a phylogenetic tree via the QIIME2 align-to-tree-mafft-fasttree pipeline, which aligns sequences with MAFFT [22] and builds a tree with FastTree [23]. The resulting feature table, taxonomy assignments, phylogenetic tree, and sample metadata were imported into R (version 4.4.0) using the phyloseq package [24] for downstream community analysis.

### Disease Assay

WT *Pst* DC3000, *hrcC* and Δ*fecA* Δ*pvdI* Δ*pvhA* were streaked on Luria-Marine (LM) medium with 100 µg/mL rifampicin and grown for two days at 28℃. Cells were resuspended in 1mL 10 mM MgCl_2_ and washed twice. Cultures were normalized to OD_600_ = 0.0002 and plated on LM+Rif100 plates to confirm that bacterial counts corresponded to 10^5^ CFU/mL. Prior to infiltration, plants were domed for several hours to increase stomata opening for ease of infiltration. Two leaves per plant were infiltrated and excess bacterial culture was blotted away. Leaves were allowed to dry, then covered and incubated at 95% humidity for three days. Leaves were weighed, surface sterilized with 5% bleach for 1 min and rinsed twice in milliQ water for 1 min each. Leaves were ground in 500 µl 10 mM MgCl_2_, serially diluted and plated on LM+Rif100 plates which were incubated at 28℃ for two days. Pathogen titers were counted and calculated as CFU/mg fresh weight. Samples were for 16S rRNA analysis were treated similarly but collected in impact resistant tubes with beads, flash frozen in liquid nitrogen and stored at -80°C until DNA extraction.

### *In planta* binary interaction assay

Bacterial strains, *S. maltophilia* (C46) and *P. chondroitinus* (C3) were streaked from glycerol stocks on R2A plates and grown at room temperature for two days. Each strain was scraped from the plates, washed twice in 1 mL 10 mM MgCl_2_, and resuspended in sterile 10 mM MgCl_2_. OD_600_ was measured and C3 and C46 were respectively diluted to a final OD_600_ of 0.1 (10^7^ CFU/mL) and 0.0125 (10^7^ CFU/mL) in sterile miliQ water. For iron treatments, 5 μM FeSO_4_ or equivalent volume of miliQ water was added to the normalized C3, C46, C3/C46, or miliQ only inoculum. Initial inoculum counts were performed for all treatments and control by plating on both R2A only and R2A with 3 µg/ml colistin sulfate (Sigma-Aldrich, CAS RN: 1264-72-8) plates to verify ∼1.0 x 10^7^ CFU/mL inoculum for each experiment. The SynCom inoculum was infiltrated into 4.5-week-old marked leaves using a 1 mL needleless syringe and allowed to dry. The plants were then covered with a dome to maintain high humidity (95%) for five days.

To quantify microbial load, inoculated leaves were removed and surface sterilized in 5% bleach for 1 min and washed twice in milliQ water for 1 min each. Leaves were placed in 2 mL impact resistant tubes containing three zirconium beads with 500 µl 10 mM MgCl_2_. Samples were ground using a tissue lyser, serially diluted, plated on R2A and/or R2A+3 µg/ml colistin sulfate plates and incubated for 2 days at room temperature. CFU/mg of fresh weight was calculated. The experiment was repeated three independent times with eight biological replicates each.

### *In vitro* binary interaction assay

To prepare 40 mM 2,2^’^-dipyridyl (DIP) solution, 93.7 mg of 2,2-dipyridyl (Goldbio, CAS RN: 366-18-7) was added to 15 mL 40% dimethyl sulfoxide (DMSO) and placed in a water bath at 68°C to dissolve. The solution was filter sterilized and stored at 4 °C in the dark.

Bacterial strains (C46, C3 and C17-2) were streaked on R2A plates and grown at room temperature (22°C) for 2 days. Each strain was scraped from plates, washed twice in 1mL 10 mM MgCl_2_, and resuspended in sterile 10 mM MgCl_2_. The target strain (C3 and C17-2) was normalized to OD_600_ of 0.375 and the attacker strain (C46) was normalized to OD_600_ of 0.5. After normalization, the attacker strain was further concentrated via resuspension in 10 mM MgCl_2_ to a fifth of its original volume. Thirty microliters of the target strain were spread on R2A + 100 uM DIP (+/-50 µM FeSo_4_) plates and allowed to dry for 10 mins. Ten microliters of the attacker strain (C46) was spotted in the center of plates and dried for an additional 20 mins. The plates were incubated at room temperature for two days and then imaged.

To quantify inhibition zones, images were loaded into ImageJ. The function “measure” was used and calibrated with the diameter of the plate (60 mm). Zone of inhibition was defined as the distance between two points at which the growth of bacterial strains are no longer visible within the same radial axis. The defined lengths were traced and measured at three different radial axes. The mean of three measurements obtained from a plate was calculated and compared with the unpaired Student’s t-test.

#### Binary Interaction between Pst DC3000 and SynCom collections

We performed binary interaction between *P. syringae Pst* DC3000 and *Pst* DC3000 Δ*fecA* Δ*pvdI* Δ*pvhA* (attacker strains) against He lab SynCom^Col-0^ and Vorholt collection (target strains). SynCom^Col-0^ and Vorholt collection were streaked on R2A and R2A+0.5% methanol and grown for two days at room temperature. Cells were scraped from plates and resuspended in 500 µl of 10 mM MgCl_2_ (OD_600_ between 0.1-0.7). Fifty microliters of target strains were spread on a 60 mm x 15 mm R2A or R2A+0.5% methanol plate and allowed to dry for 5-10 mins. WT *Pst* DC3000 and *Pst* DC3000 Δ*fecA* Δ*pvdI* Δ*pvhA* were normalized to OD_600_ 1.5 (∼3 x 10^9^ CFU/mL) and 5 ul spotted onto the center of the plate and allowed to dry. Plates were incubated for 2 days at room temperature and imaged. Zones of inhibition were determined as described above.

### Chrome Azurol S (CAS) assay

The original CAS assay was developed by Schwyn and Neilands [25].

Bacterial strains were streaked on R2A plates and grown at room temperature (22 °C) for 2 days. Each strain was scraped from plates, washed twice in 1mL 10 mM MgCl_2_, and resuspended in sterile 10 mM MgCl_2_.

#### Solution Preparation

To prepare shuttle solution, 5-sulfosalicylic acid dyhydrate (Sigma Aldrich, CAS RN: 5965-83-3) was added to miliQ water to the final concentration of 0.2 M and stored in the dark. To prepare PIPES buffer, 15.1 g of PIPES (Sigma-Aldrich, CAS RN: 5625-37-6) was dissolved in 80 mL of MiliQ water and pH was adjusted to 6.8 with 5 M NaOH. The final volume was adjusted to 100 mL with milIQ water. CAS blue dye was prepared as previously described [26]. To prepare CAS assay solution, 50 mL of miliQ water, 10 mL of PIPES buffer, and 10 mL of CAS blue dye were added and mixed.

#### Liquid CAS

For liquid CAS assays, bacterial strains were cultured in R2A liquid medium containing 100, 200, 400 μM of DIP or equivalent concentration of DMSO for 3 days at 28 °C. Absorbance at OD_600_ was measured for each bacterial strain to account for the impact of DIP on growth and 1 mL of culture from each treatment was transferred to eppendorf tubes and centrifuged for 30 mins at 5,700 rpm at 4 °C. The supernatant was removed and filtered through a 0.22 μm filter and then transferred to a fresh eppendorf tube. The filtered supernatant (500 µl) was added to 500 μL of CAS solution and 10 μL of shuttle solution and mixed by gently pipetting up and down. The mixture was incubated at room temperature for 24 h and absorbance at OD_630_ was measured.

For plate assays, we used the overlay method [27]. Thirty microliters of washed bacterial strain (OD_600_ between 0.3 and 1.0) was spotted on R2A plates containing 200 µM DIP and incubated at 28°C for 3 days. Five milliliters of CAS overlay medium (0.7% agar, 10 mL CAS blue dye, 10 mL PIPES) was poured onto culture plates and incubated at room temperature for 24 h and subsequently imaged.

### Western blot

Roots from 4.5-week-old Arabidopsis thaliana Col-0 WT and *mfec* plants were collected, cleaned and weighed before placing in 2 mL impact resistant tubes containing beads. Similar mass of roots were used across all genotypes. Root samples were flash frozen in liquid nitrogen and homogenized using a tissue lyser and boiled for 5 mins at 95°C in 1x LDS sample buffer (Genscript) with 2.5% β-mercaptoethanol. Samples were centrifuged and the supernatant transferred to a new tube. Samples were ran on a 4-12% SURE PAGE gel (Genscript) for ∼50 mins at 150 V and protein transfer performed by wet transfer for 1h at 30V. Blots were blocked in 5% skim milk in 1x TBST (Tris-buffered saline with 0.05% Tween-20) for 1h at room temperature, then incubated with primary antibody (α-IRT1, Agrisera; 1:3000) overnight at 4°C, followed by secondary antibody (goat α-rabbit IgG Agrisera; 1:5000) for 1h at room temperature. The blot was imaged using the iBright 1500 (Invitrogen). Amido black staining was performed on the same membrane to visualize total protein as a loading control.

For Western blot of AHA2 and BAK1, two leaf punches per leaf were collected from two leaves of 4-week-old Arabidopsis Col-0 WT and *mfec* plants. Punches were homogenized in 200 µl 1x RIPA buffer using a tissue lyser. Lysates were incubated at 4°C for 30 min prior to centrifugation at 10,000 x g to remove cell debris. Protein concentration was determined using a Bradford assay and samples were normalized to 2 µg total protein per lane. Samples were resolved on a 10% SDS-PAGE gel for 50-60 min at 150 V followed by semi-dry transfer to a membrane. The membrane was cut into two strips and probed separately for AHA2 and BAK1. Blots were blocked in 5% skim milk in 1x TBST for 1 h at room temperature, then incubated with either α-AHA2 (Agrisera; 1:1000) or α-BAK1 (Agrisera; 1:5000) primary antibodies at 4°C overnight. Blots were incubated with secondary antibody (goat α-rabbit IgG, Agrisera; 1:5000) for 1h at room temperature. Blots were imaged using an iBright 1500. BAK1 served as the membrane loading control.

### Growth Curve

WT *Pst* DC3000 and *Pst* DC3000 Δ*fecA* Δ*pvdI* Δ*pvhA* were streaked on LM+Rif100 µg/mL plates and grown for two days at 28°C. Cells were resuspended in 1 mL of LM medium, washed twice and normalized to a starting OD_600_ of 0.05 in R2A. A kinetic growth assay was performed in a flat-bottom 96-well plate, and OD_600_ was measured at 28°C every 40 mins for 24h using a microplate reader (SpectraMax M2) with continuous shaking between measurements.

### Inductively Coupled Mass Spectrometry (ICP-MS)

Leaves from 4-week-old WT Col-0 and *mfec* plants were harvested and weighed (∼1g of fresh weight is approximately 100 mg dry weight). Leaves were placed in grease bags and dried in an oven at 65°C for two days. Dried leaf tissue was digested by microwave digestion (Mars6, CEM Microwave) in Xpress vessels containing in 4.5 mL HNO_3_ and 1.5 mL H_2_O_2_. The exact sample mass transferred to the vessels was recorded. Digestion was performed using a preset plant tissue program (30 min digestion, 20 min cool-down). All digestions included ∼0.1g tomato leaf NIST standard (SKU: 1573a) reference material.

### HPTS pH Imaging

*Arabidopsis* WT and *mfec* plants were grown as described above for 3.5 weeks. Before sampling, all plants were domed to keep at high relative humidity (∼95%) for 24 h. Leaf discs of 4mm were made and immediately put into 2 mM 8-hydroxypyrene-1,3,6-trisulfonic acid trisodium (HPTS) with 0.005% silwet-77 (Plant Media) solution. A pressure of -0.08 MPa was applied to the samples for 2 mins and then released slowly for ∼40 s to ambient pressure. Samples were swirled occasionally between vacuum infiltration. All the samples were covered with foil and equilibrated at room temperature for at least 1h before imaging.

Leaf discs were mounted on a slide in HPTS staining solution and covered with a coverslip. The abaxial side was imaged for apoplastic pH using an inverted Zeiss 880 confocal microscope. Fluorescent signals and calculation method were previously described [28]. The relative pH was determined by the value ratio of channel 405nm/458nm. Higher values mean more acidic relative to WT. Experiments include four plants each representing a biological replicate.

### FeGenie Analyisis

To identify iron-related genes across members of SynCom^Col-0^ and C46, GenBank-formatted genome assembly files for all 49 strains were analyzed using FeGenie [29], a bioinformatics tool that screens genome and metagenome assemblies against a curated database of hidden Markov models (HMMs) covering genes involved in iron acquisition, storage, transport, and redox-cycling. For each genome, the percentage of protein-coding genes (ORFs) assigned to each iron-related functional category was calculated as gene count for that category divided by total ORF count, following FeGenie’s standard normalization. Genomes were grouped by phylum and hierarchically clustered based on these values to generate the heatmap shown in Fig. 4d.

### Statistical Analysis

All data points on the graphs represent biological replicates. All experiments were performed at least three independent times. Graphs and statistics were generated using GraphPad Prism v11.0.2. Data are presented as +/-SEM. Samples with more than two experimental groups were analyzed using one-wat ANOVA with Dunnet’s multiple comparison test. An unpaired two-tailed Student’s t-test was used for experiments with two experimental groups as shown in Fig. 4e. Different letters indicate statistically significant groups. No statistical methods were used to predetermine sample size. The experiments were not randomized and experiments were performed without investigator blinding.

## AUTHOR CONTRIBUTIONS

S.S.W. conceived, designed, performed, analyzed, interpreted experiments and wrote the manuscript. J.X. analyzed the metatranscriptomics data and T.C. performed *in planta* metatranscriptome experiments in WT and *mfec* plants. M.K. performed *in vitro* experiments, supplementation experiments with supervision and help from S.S.W. M.K. helped with preparing samples for 16S analysis. J.Z. performed HPTS experiments. S.Y.H. conceived, interpreted data, supervised and co-wrote the manuscript. All authors have read the manuscript.

## ACKNOWLEDGEMENTS

We thank Hongze Wang from Jian-Min Zhou’s lab at Yazhouwan National Laboratory for sharing an initial protocol of HPTS staining in *Arabidopsis* leaves. We thank the Duke Phytotron staff for technical assistance with preparing soil and maintaining plant growth chambers. We are also grateful to members of the He lab for insights on the project and critically reading the manuscript. Also, thanks to undergraduate Laya Tummala for help during the initial stages of the project. This work was supported by the HHMI Hanna Gray Fellowship awarded to S.S.W. and M.K. is supported by HHMI Cech Fellowship and Duke University Undergraduate Research Student Assistantship award. Support for this work is also provided by the National Institutes of Allergy and Infectious Diseases (grant AI15544) and Duke Science and Technology Initiative awarded to S.Y.H. S.Y.H. is an Investigator at Howard Hughes Medical Institute. The 16S and RNA-sequencing data presented herein were acquired, in part, using instrumentation in the Genomics Core (RRID:SCR_012406), supported by Michigan State University’s Office of Research & Innovation.

## DATA AND CODE AVAILABILITY

Raw metatranscriptomic sequencing reads from C46 alone and C46 in SynCom experiments are available in the NCBI Sequence Read Archive (SRA) under BioProject PRJNA864025. RNA-seq raw sequencing reads and processed data have been deposited in the NCBI Gene Expression Omnibus (GEO) database under accession GSE345708. Raw 16S rRNA gene sequences are available in the NCBI Sequence Read Archive (SRA) under BioProject PRJNA1521988 (accession numbers SAMN62822731-SAMN62822826, iron treatment experiment) and BioProject PRJNA1522051 (accession numbers SAMN62825832-SAMN62825855, *Pst* infection experiment). 16S rRNA gene sequences were processed using QIIME2 (version 2025.7), with taxonomy assigned using a naive Bayes classifier trained on the SILVA 138.2 SSU NR99. No new code was generated in this manuscript.

## SOURCE DATA

All source data will be included in the final manuscript.

