## Supplementary figures and images for "Iron as a principal mediator of dysbiosis prevention in the phyllosphere"

### Supplementary Western blot images

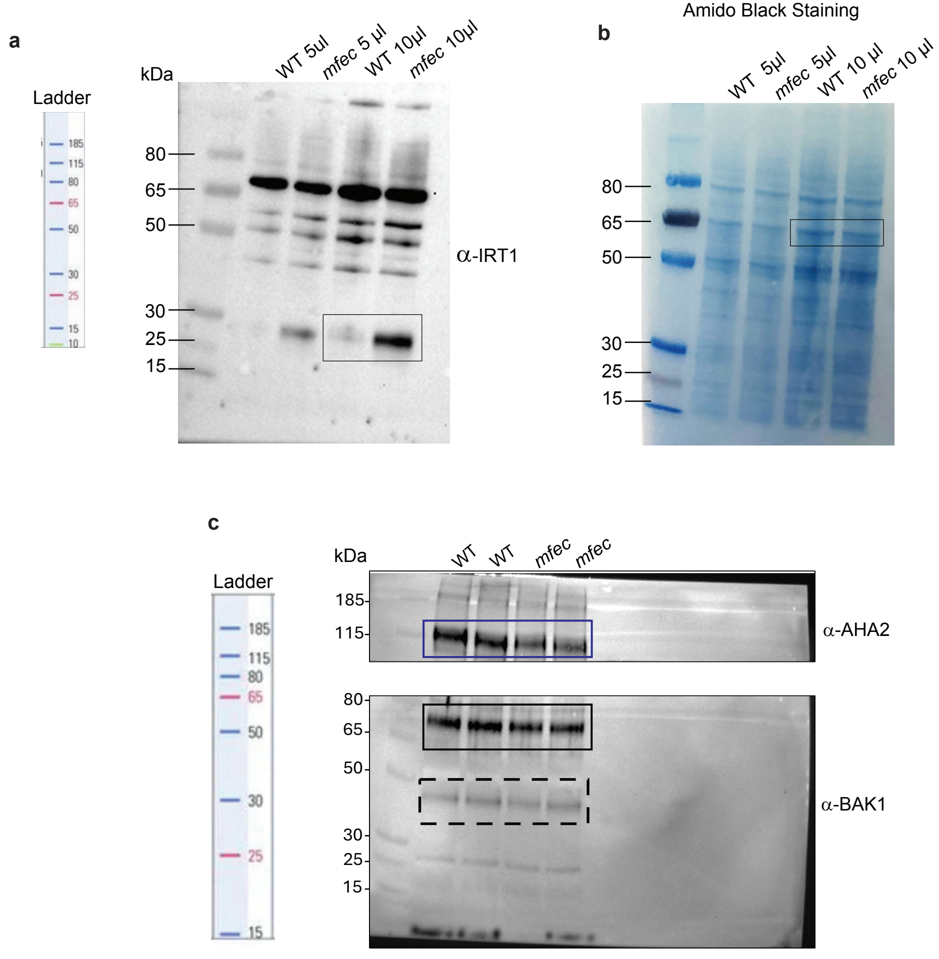


Supplementary Figure 1
